# Infralimbic cortical glutamate output is necessary for the neural and behavioral consequences of chronic stress

**DOI:** 10.1101/2020.06.30.181115

**Authors:** Sebastian A. Pace, Connor Christensen, Morgan K. Schackmuth, Tyler Wallace, Jessica M. McKlveen, Will Beischel, Rachel Morano, Jessie R. Scheimann, Steven P. Wilson, James P. Herman, Brent Myers

## Abstract

Exposure to prolonged stress is a major risk-factor for psychiatric disorders such as generalized anxiety and major depressive disorder (MDD). Human imaging studies have identified structural and functional abnormalities in the prefrontal cortex of MDD patients, particularly Brodmann’s area 25 (BA25). Further, deep brain stimulation of BA25 reduces symptoms of treatment-resistant depression. The rat homolog of BA25 is the infralimbic cortex (IL), which is critical for cognitive appraisal, executive function, and physiological stress reactivity. Previous studies indicate that the IL undergoes stress-induced changes in excitatory/inhibitory balance culminating in reduced activity of glutamate output neurons. However, the regulatory role of IL glutamate output in mood-related behaviors after chronic variable stress (CVS) is unknown. Here, we utilized a lentiviral-packaged small-interfering RNA to reduce translation of vesicular glutamate transporter 1 (vGluT1 siRNA), thereby constraining IL glutamate output. This viral-mediated gene transfer was used in conjunction with a quantitative anatomical analysis of cells expressing the stable immediate-early gene product ΔFosB, which accumulates in response to repeated neural activation. Through assessment of ΔFosB-expressing neurons across the frontal lobe in adult male rats, we mapped regions altered by chronic stress and determined the coordinating role of the IL in frontal cortical plasticity. Specifically, CVS-exposed rats had increased density of ΔFosB-expressing cells in the IL and decreased density in the anterior insula. The latter effect was dependent on IL glutamate output. Next, we examined the interaction of CVS and reduced IL glutamate output in behavioral assays examining coping, anxiety-like behavior, associative learning, and nociception. IL glutamate knockdown decreased immobility during the forced swim test compared to GFP controls, both in rats exposed to CVS as well as rats without previous stress exposure. Further, vGluT1 siRNA prevented CVS-induced avoidance behaviors, while also reducing risk aversion and passive coping. Ultimately, this study identifies the necessity of IL glutamatergic output for regulating frontal cortical neural activity and behavior following chronic stress. These findings also highlight how disruption of excitatory/inhibitory balance within specific frontal cortical cell populations may impact neurobehavioral adaptation and lead to stress-related disorders.

**Highlights:** Chronic stress increased ΔFosB in the infralimbic cortex and decreased insular ΔFosB

Decreased insular ΔFosB was dependent on infralimbic glutamate output

Knockdown of infralimbic glutamate release reduced passive coping

Avoidance behaviors after chronic stress were dependent on infralimbic glutamate

Infralimbic projections innervated excitatory and inhibitory neurons in the insula

## 1. Introduction

The medial prefrontal cortex (mPFC) contains multiple cell groups instrumental for cognition, executive function, and emotion (Bechara et al., 2000; Drevets et al., 1997). Additionally, the mPFC regulates the activity of autonomic, endocrine, and behavioral systems to facilitate appraisal and contextual adaptation (Duncan, 2001; McKlveen et al., 2015; Myers, 2017; Ulrich-Lai & Herman, 2009). Consequently, alterations in the structure and function of the mPFC associate with behaviors such as social avoidance, despair, and fear, potentially contributing to psychiatric conditions such as major depressive disorder (MDD), generalized anxiety disorder, and post-traumatic stress disorder (Arnsten, 2009; Covington et al., 2010; Drevets et al., 2008a; Drevets et al., 1997; Holmes et al., 2018; Liotti et al., 2000; Mayberg et al., 1999; Schiller et al., 2008; Warden et al., 2012). More specifically, neuroimaging studies implicate the subgenual region, Brodmann’s area 25 (BA25), in the pathology of depression (Drevets et al., 2008a; Drevets et al., 1997; Drevets et al., 2008b). In MDD patients, BA25 has decreased glucose metabolism and reduced postmortem gray matter (Drevets et al., 2008); furthermore, deep brain stimulation of BA25 has been used as a therapeutic intervention for treatment-resistant depression (Kennedy et al., 2011; Lozano et al., 2008; Mayberg, 2005).

Comparative analyses indicate that the rodent mPFC contains topographically-distinct regions homologous to the human cortex (Ongür & Price, 2000; Uylings et al., 2003). The BA25 homolog in rodents, infralimbic cortex (IL), regulates neuroendocrine and cardiovascular responses to chronic stress (McKlveen et al., 2013; Myers et al., 2017; Schaeuble et al., 2019). Moreover, chronic stress leads to dendritic retraction and increased inhibition of glutamate-releasing IL pyramidal neurons, facilitating excitatory/inhibitory imbalance (Anderson et al., 2019; Cook & Wellman, 2004; Goldwater et al., 2009; McKlveen et al., 2016; McKlveen et al., 2019; Radley et al., 2004). Chronic stress also alters mPFC immediate-early gene expression, including Arc, Zif268, and FosB/ΔFosB (Covington et al., 2010; Flak et al., 2012; Vialou et al., 2015). Within the Fos transcription factor family, FosB and ΔFosB are splice variants of the *fosb* gene (McClung et al., 2004; Nakabeppu & Nathans, 1991). FosB is expressed transiently after a stimulus, while the stable truncated protein ΔFosB accumulates in response to repeated stimulation (McClung et al., 2004; Nestler et al., 2002). Thus, ΔFosB accumulation identifies chronically-activated neurons and serves as a molecular marker for cellular plasticity in response to long-term stimulation (Flak et al., 2012; Nestler, 2015). Behaviors induced by chronic stress, including passive coping and avoidance, likely arise from cellular- and circuit-level changes in neural activity; therefore, ΔFosB expression in distinct frontal cortical cell populations may represent a unique neural signature for chronic stress.

In the current study, high-resolution quantitative analysis of ΔFosB-expressing neurons throughout the prefrontal, orbital, and insular cortices queried the role of IL glutamate release in frontal cortical activation after chronic stress. An interfering RNA approach was used to target the expression of IL vesicular glutamate transporter 1 (vGluT1) and reduce glutamate outflow. vGluT1 is essential for pre-synaptic vesicular glutamate packaging, release, and synaptic transmission in pyramidal glutamatergic neurons (Schuske & Jorgensen, 2004; Wojcik et al., 2004; Ziegler et al., 2002); furthermore, vGluT1 knockdown selectively reduces vGluT1 mRNA and protein expression (Myers et al., 2017). This approach was also used to examine the importance of IL glutamate output for the behavioral consequences of chronic variable stress (CVS), including passive coping and avoidance. Ultimately, these studies identify a region-specific cellular basis for adaptation to chronic stress and suggest that circuit-level mechanisms in the frontal cortex may account for stress-related disorders.

## 2. Methods

### 2.1. Animals

Adult male Sprague-Dawley rats from Envigo (Indianapolis, IN or Denver, CO) weighing 250-300 g were single-housed in a temperature and humidity-controlled vivarium with a 12-hour light-dark cycle (lights on at 0600, and off at 1800). Rats were acclimated to the vivarium for 1 week before the start of the experiment. Water and chow were available *ad libitum* throughout the experiment. All procedures and protocols were approved by the Institutional Animal Care and Use Committee of either the University of Cincinnati (protocol: 04-08-03-01) or Colorado State University (protocol: 16-6871A) and complied with the National Institutes of Health Guidelines for the Care and Use of Laboratory Animals. The cumulative sequence of procedures used in the current experiments received veterinary consultation and all animals had daily welfare assessments by veterinary and/or animal medical service staff.

### 2.2. Lentiviral construct

An antisense RNA approach was used to knockdown vGluT1 as previously described (Myers et al., 2017; Schaeuble et al., 2019). Briefly, a lentivirus transfer vector was constructed from a third-generation self-inactivating transfer vector (de Almeida et al., 2001). The transfer virus was transfected with packaging plasmids, psPAX2, pRSV-Rev, and pMD2.G (Addgene, Cambridge, MA) in 293T cells. Viruses underwent high-speed centrifugation for concentration, centrifugation through 20% sucrose/Dulbecco’s phosphate-buffered saline for purification and were stored at –80 C° in 10% sucrose/Dulbecco’s phosphate-buffered saline. Quantitative real-time polymerase chain reaction determined virus particle concentrations for proviral DNA 24-hours following transduction in 293T cells and was shown as transducing units per microliter (tu/µL). Rat vGluT1 complementary DNA synthesized included 151 bp of the 3’ coding region and 212 bp of the 3’ untranslated region consistent to GenBank accession no. NM_053859 nucleotides 1656 to 2018. This region circumvents common glutamate transporter transmembrane domains and shares little homology with vGluT2 and vGluT3. The region was cloned in an antisense manner into a lentivirus transfer vector that expressed an enhanced green fluorescent protein (eGFP) reporter. This vector is under the control of the phosphoglycerate kinase-1 promoter, which expresses preferentially to rat brain neurons (Grillo et al., 2007; Grillo et al., 2015; Krause et al., 2011; Myers et al., 2017; Wood et al., 2019). Additionally, a control virus containing a transfer vector with a phosphoglycerate kinase-1 promoter drove expression of eGFP.

### 2.3. Stereotaxic surgery

Rats were anesthetized via intraperitoneal injection of 90 mg/kg ketamine and 10 mg/kg xylazine and given analgesic (2 mg/kg butorphanol, subcutaneous) and antibiotic (5 mg/kg gentamicin, intramuscular). Rats received 1 μL (5 × 10^6^ tu/µL titer) bilateral microinjections of siRNA-expressing or GFP-expressing virus into the IL (2.9 mm rostral to bregma, 0.6 mm lateral to midline, and 4.2 mm ventral from dura), as previously described (Myers et al., 2017). Microinjections were made with a 25-gauge, 2-μL microsyringe (Hamilton, Reno, NV) attached to a microinjector (Kopf, Tujunga, CA) to infuse virus at a rate of 5 min/µL. The microsyringe was left in place for 5 min before and after injections to allow for diffusion of the virus and reduce tissue damage. Post-injection, rats recovered for 6 weeks before further experimental procedures, corresponding to timeframes from prior studies using lentivirus (Myers et al., 2017; Schaeuble et al., 2019; Wood et al., 2019).

### 2.4. Chronic variable stress

CVS was comprised of twice daily (AM and PM) repeated and unpredictable stressors presented in a randomized manner, including exposure to a cold room (4°C, 1 hour), shaker stress (100 rpm, 1 hour), restraint (Plexiglas tube, 30 minutes), and hypoxia (8% oxygen, 30 minutes) (Flak et al., 2014; Ghosal et al., 2014). As described below, behavioral tests were included as stressors at the beginning and end of CVS, including a brightly-lit open field (1 m^2^, 5 minutes), forced swim (23–27°C, 10 minutes), and elevated plus maze (0.5 m, 5 minutes). Additionally, overnight stressors were variably included, consisting of wet bedding (16 hours), social crowding (6–8 rats/cage, 16 hours), and restricted housing (mouse cage, 16 hours).

### 2.5. Experimental design

The data in the current report were collected from three separate experiments. For experiment 1 (Fig.1A), rats were divided into three groups: 1) Rats injected with a GFP-expressing construct and remaining unstressed (No CVS GFP, n = 11), 2) Rats with GFP injections that were exposed to CVS (CVS GFP, n = 14), or 3) CVS-exposed rats with siRNA treatment (CVS siRNA, n = 14). Behavioral assays were added to the CVS protocol. As the AM stressor of CVS day 1, previously unstressed CVS GFP and CVS siRNA rats underwent the open field test with novel object interaction. On the morning of day 2, the CVS groups completed a shock probe defensive burying assay. On the mornings of CVS days 12 and 13, the CVS groups were re-tested in the shock probe assay, open field, and novel object interaction. On the final morning of CVS, both CVS groups underwent the forced swim test. Animals were then allowed 24 hours of recovery to allow resolution of acute stress-induced FosB expression and more selectively isolate ΔFosB expression (McClung et al., 2004; Vialou et al., 2015). All 3 groups were then euthanized for tissue collection as described below.

**Figure 1:**
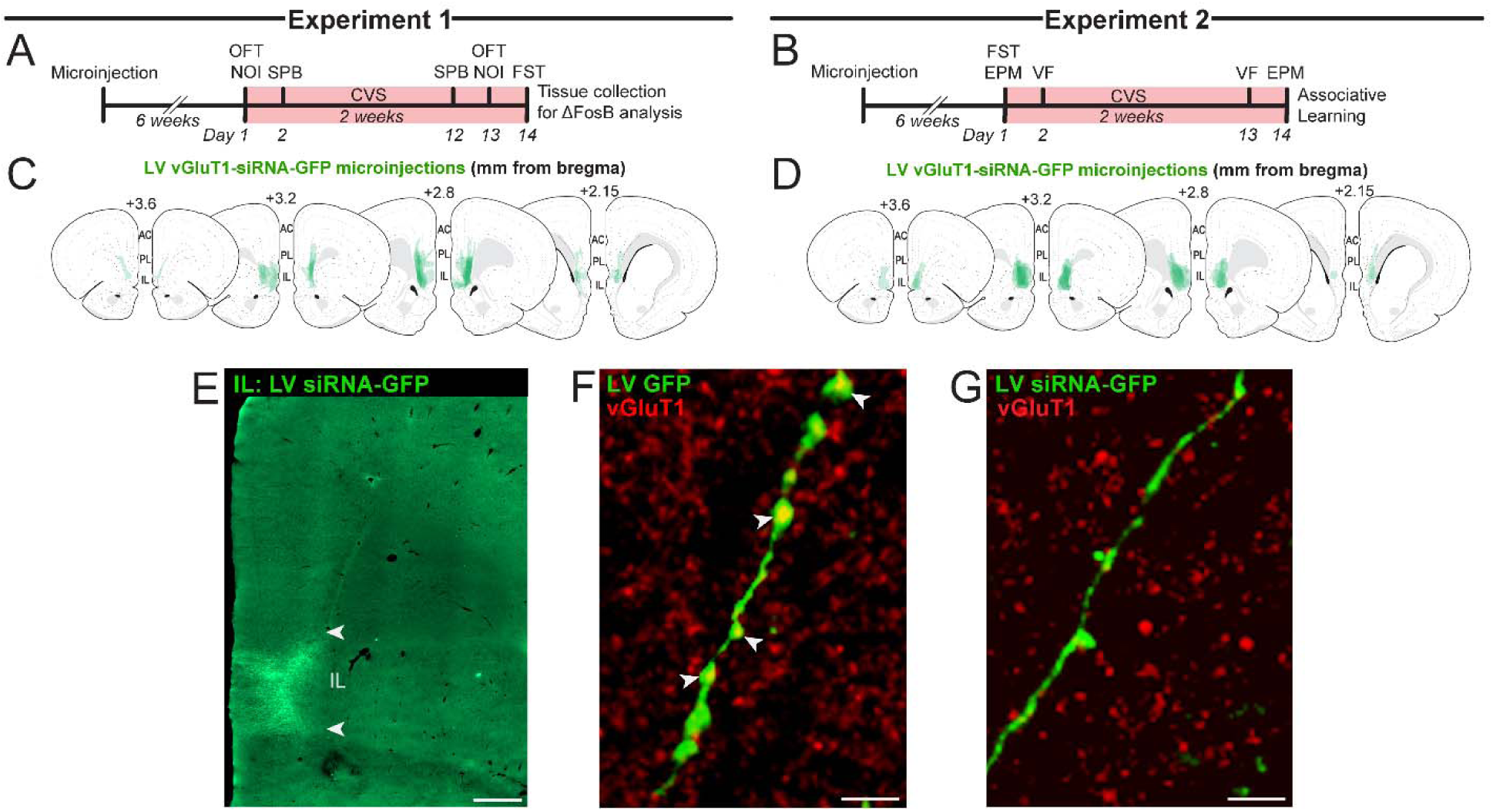
Experimental timelines and design. Timelines for experiments 1 (A) and 2 (B). Lentiviral vGluT1 siRNA-GFP microinjections were mapped onto Swanson Rat Brain Atlas (3^rd^ edition) coronal sections (C, D). Injections targeted the IL, scale bar: 500 μm (E). In the GFP control group, vGluT1 protein was localized to GFP-labeled projections (F). Co-localization was reduced by siRNA treatment, scale bars: 5 μm (G). AC: anterior cingulate cortex, CVS: chronic variable stress, EPM: elevated plus maze, FST: forced swim test, GFP: green fluorescent protein, IL: infralimbic cortex, LV: lentivirus, NOI: novel object interaction, OFT: open field test, PL: prelimbic cortex, siRNA: small interfering RNA, SPB: shock probe burying, VF: Von Frey test, vGluT1: vesicular glutamate transporter 1.

The second experiment (Fig.1B) used a similar 3-group design [No CVS GFP (n = 8), CVS GFP (n = 10), and CVS siRNA (n = 12)]. In this cohort, the forced swim test was the AM stressor of CVS day 1 followed by elevated plus maze in the PM. On Day 2, the Von Frey method was used to determine nociceptive thresholds in CVS GFP and CVS siRNA rats. On days 13 and 14, both CVS groups repeated the Von Frey assessment and elevated plus maze, respectively. After the completion of CVS, all 3 groups received tone-shock conditioning on day 15. In a third experiment, an AAV-packaged construct was injected in the IL (n = 4) to permit genetically-encoded identification of IL pre-synaptic terminals in the insula. For all experiments, treatment assignments were randomized and all experimenters blinded.

### 2.6. Forced swim test

The forced swim test is commonly used to assess coping behavior (Molendijk & de Kloet, 2019); furthermore, antidepressants increase active coping and reduce immobility (Smith et al., 2017, 2018; Solomon et al., 2014). A modified test was used that involves a single swim exposure (Cryan et al., 2005), which differs from the original Porsolt forced swim test that requires repeated exposure to examine learned helplessness (Porsolt et al., 1977). Rats were placed in an open-top cylinder (61 cm in height × 19 cm in diameter) filled with 40 cm of 25 +/-2 °C water for 10 min. Behavior was recorded with an overhead mounted video camera. The video was analyzed by a blinded observer every 5 s to score immobility or activity (swimming, climbing, and diving).

### 2.7. Open field test/novel object interaction

The combined open field test/novel object interaction was used as previously described (Myers et al., 2016) to assess general locomotor activity, approach/avoidance, and novelty preference/aversion (Belzung & Griebel, 2001). Rats were placed in a black acrylic 1 m^2^ square field with 30 cm tall walls for 5 min. Clever TopScan behavioral analysis software determined total distance traveled and time spent in the center of the field (central 0.5 m square). For novel object interaction, a novel object was placed in the center of the field with the surrounding 10 cm^2^ area defined as the interaction zone. Time in the interaction zone and object sniffing were quantified over 5 min.

### 2.8. Elevated plus maze

The elevated plus maze examines anxiety-like behaviors based on time spent exploring open spaces (Pellow et al., 1985). Under low light conditions, rats were placed on an open arm facing the center of an elevated (0.5 m) platform composed of 2 open arms and 2 enclosed arms (Myers et al., 2016). Clever TopScan analysis software quantified time spent in open and closed arms, as well as the exploration of the outer end of open arms (most distal 30 cm).

### 2.9. Shock probe defensive burying

The shock probe burying test was used to characterize defensive behavior and coping style (Boersma et al., 2014). Rats were placed in a novel shoebox cage with clean bedding for 10 min of habituation. Next, an electric prod (8.5 cm long, 1 cm diameter) was placed through a small opening in the front of the cage. A 2.5 mA shock was delivered when the experimental rat contacted the prod. Over a 5 min testing period, probe contacts and stretch-attend behaviors toward the probe were determined by a treatment-blind observer. After the first contact, time spent burying the probe or freezing was also quantified.

### 2.10. Von Frey assay

Von Frey filaments were used to determine cutaneous sensitivity to noxious stimuli (Myers et al., 2007). Rats were placed in a clear Plexiglass container (21 cm × 27 cm × 15 cm) on an elevated (45 cm) mesh floor (12 mm × 12 mm grid) for 30 min of habituation. Von Frey hairs (Stoetling, Wood Dale, IL) were pressed perpendicularly against the plantar region of the hind paw at increasing forces. The force eliciting reflexive withdrawal of the hind paw was determined three times with the values averaged for each animal.

### 2.11. Associative learning

To determine whether vGluT1 knockdown might impact associative learning after chronic stress, we used an abbreviated version of a tone-shock fear conditioning protocol (Mueller et al., & Quirk, 2011; Vollmer et al., 2016). Auditory conditioning was carried out in sound-attenuating chambers with stainless-steel bar flooring (Med Associates, Burlington, VT). Two tones (70 dB, 2 kHz, 30 s with a 3 min intertrial interval) were paired with footshocks (0.5 mA, 0.5 s) followed by a third tone without shock. Learned acquisition of freezing behavior during tone presentation was measured as time spent freezing during the 30 s tone (FreezeScan, Clever Systems, Reston, VA).

### 2.12. SynaptoTag

To investigate IL projections to the insula, rats were prepared for stereotaxic surgery as described above. In this third experiment, 50 nL an adeno-associated virus (AAV) was injected unilaterally in the IL (4.7 × 10^8^ tu/μl). The AAV (Stanford Gene Vector and Virus Core, Palo Alto, CA) carried a construct under the control of the human synapsin promoter (hSyn1) that codes for mCherry expression in neuronal soma and axons, as well as enhanced GFP conjugated to synaptobrevin-2. Thus, neurons expressing this construct are marked by mCherry expression with pre-synaptic terminals reported by GFP (Xu & Südhof, 2013). Following injections, rats recovered for 6 weeks to allow reporter expression before tissue collection.

### 2.13. Tissue collection

After experiments, all animals were euthanized with sodium pentobarbital (≥ 100 mg/kg, intraperitoneal) and transcardially perfused with 0.9% saline followed by 4% phosphate-buffered paraformaldehyde. Brains were post-fixed in paraformaldehyde for 24 hours and then stored in 30% sucrose at 4°C. Brains were subsequently sectioned (30 μm thick 1:12 serial coronal sections) and stored in cryoprotectant solution at -20 °C until immunohistochemistry.

### 2.14. Immunohistochemistry

To determine injection placement, GFP labeling from vGluT1-siRNA and GFP injections was used to map the extent of spread. Tissue sections were washed in phosphate-buffered saline (PBS) (5 × 5 min), incubated in blocking solution (PBS, 0.1% bovine serum albumin, and 0.2% TritonX-100) for an hour, then placed overnight at 4 ^0^C in rabbit anti-GFP primary antibody (1:1000 in blocking solution, Invitrogen, La Jolla, CA). Following incubation, tissue sections were washed and placed into Alexa 488-conjugated donkey anti-rabbit immunoglobulin G (IgG; 1:500 in blocking solution; Jackson Immunoresearch, West Grove, PA) for 30 minutes. Additionally, dual-fluorescent immunolabeling of GFP and vGluT1 was carried out to visualize knockdown. After GFP labeling, vGluT1 was visualized with rabbit anti-vGluT1 primary antibody (1:1000 in blocking solution; Synaptic Systems, Goettingen, Germany) and Cy3-conjugated donkey anti-rabbit IgG (1:500 in PBS; Jackson Immunoresearch, West Grove, PA). After a final wash (5 × 5 min), tissue was mounted with polyvinyl alcohol mounting medium (MilliporeSigma, Burlington, MA) and coverslipped (Thermo Scientific, Portsmouth NH).

For chromogen labeling of FosB/ΔFosB, coronal brain sections were removed from cryoprotectant and rinsed in PBS (5 × 5 min) at room temperature. Sections were incubated in 0.03% hydrogen peroxide for 10 min, rinsed, and placed in blocking solution (PBS, 0.1% bovine serum albumin, and 0.2% Triton X-100) for 1 hour. Next, sections were incubated overnight in rabbit anti-FosB/ΔFosB primary antibody (1:300 in blocking solution, H75, Santa Cruz Biotechnologies, Santa Cruz, CA). The antibody was raised against amino acids 75-100 of human FosB and detects two bands by western blot, a cleaved form at 35-37 kDa (ΔFosB) and an uncleaved form at 45 kDa (FosB) (Marttila et al, 2006). Expression of uncleaved FosB returns to baseline within 6 hours of stimulus onset (Nestler et al, 2001); therefore, immunoreactivity predominately represents ΔFosB expression as euthanasia and fixation occurred 24 hours after the final stressor. Following overnight incubation in primary antibody, sections were rinsed in PBS (5 × 5 min) and incubated in biotinylated anti-rabbit secondary antibody (1:1000 in PBS with 0.1% bovine serum albumin, Vector Laboratories, Inc., Burlingame, CA) for 1 hour. Sections were then rinsed with PBS (5 × 5 min) before placement in Vectastain ABC Solution (1:1000; Vector Laboratories) for 1 hour prior to rinsing (5 × 5 min) and incubating in diaminobenzidine and hydrogen peroxide (0.02% diaminobenzidine and 0.09% hydrogen peroxide in PBS) for 10 min. Sections were rinsed in PBS, mounted, dehydrated through increasing concentrations of ethanol, and coverslipped with DPX (Sigma Aldrich).

For FosB/ΔFosB dual-fluorescent immunolabeling, tissue was washed (5 × 5 min) and placed in a low detergent blocking solution (PBS, 0.1% bovine serum albumin, and 0.025% Triton X-100) at room temperature. The tissue was then incubated overnight in rabbit anti-ΔFosB (1:2000 in blocking solution, Cell Signaling, Danvers, MA) overnight. The original anti-FosB/ΔFosB was unavailable as the manufacturer stopped the production of the antibody. Nevertheless, western blot analysis from the manufacturer demonstrates both antibodies label a similar 35-kDa form of ΔFosB. The next day, tissue was washed (5 × 5 min) and incubated in Alexa 488 goat anti-rabbit at 1:1000 in PBS for 30 min. Following FosB/ΔFosB fluorescent immunolabeling, either Ca^2+^/calmodulin-dependent protein kinase II alpha (CaMKIIα) or glutamic acid decarboxylase, 67 kDa isoform (GAD67) were labeled to determine the neurochemistry of FosB/ΔFosB-expressing cells. CaMKIIα is widely used as a marker for pyramidal glutamate projection neurons in the cortex (Liu & Jones, 1996) while GAD67 is used to distinguish local inhibitory interneurons (McGeer & McGeer, 1975). For CaMKIIα immunofluorescence, tissue underwent an extended wash in PBS (5 × 10 min) and was blocked again in a detergent-free solution (PBS, 0.1% bovine serum albumin) for 1 hour. Next, the tissue was incubated overnight in mouse polyclonal anti-CaMKIIα (1:200 in detergent-free blocking solution, GeneTex, Irvine, CA). The next day, the tissue was washed (5 × 5 min) and incubated in Cy3-conjugated donkey anti-mouse (1:500 in PBS; Jackson Immunoresearch, West Grove, PA) for 30 min. A final wash (5 × 5 min) preceded tissue mounting with polyvinyl alcohol mounting medium (MilliporeSigma, Burlington, MA), coverslipping (Thermo Scientific, Portsmouth NH), and imaging of FosB/ΔFosB and CaMKIIα double-labeled tissue. To label GAD67, tissue was washed (5 × 5 min) following FosB/ΔFosB labeling and incubated in blocking solution (PBS, 0.1% bovine serum albumin, and 0.2% TritonX-100) for 1 hour. Next, tissue was placed in mouse monoclonal anti-GAD67 (MilliporeSigma, Burlington, MA) at 1:1000 in PBS overnight. The following day, tissue was washed (5 × 5 min) and placed in Cy3 donkey anti-mouse at 1:500 in PBS for 30 min. Finally, the tissue was washed (5 × 5 min), mounted with polyvinyl medium, and coverslipped for imaging ΔFosB and GAD67 dual immunofluorescence.

For experiment 3, double labeling identified the neuronal phenotype of IL-targeted insular cortex neurons. The genetically-encoded synaptobrevin-GFP was amplified with anti-GFP immunohistochemistry followed immunolabeling for CaMKIIα or GAD67. For GFP immunohistochemistry, tissue was washed (5 × 5 min), incubated in blocking solution for 1 hour then placed in rabbit anti-GFP primary as described above. Following incubation, tissue sections were washed and placed into Alexa 488-conjugated donkey anti-rabbit immunoglobulin G for 30 minutes. Subsequently, to identify IL projections to insular excitatory neurons, CaMKIIα was incubated in mouse polyclonal anti-CaMKIIα overnight as described above. The next day, the tissue was washed (5 × 5 min) and incubated in Cy3 donkey anti-mouse for 30 min. To examine potential IL circuits targeting insular interneurons, GFP-labeled tissue was washed (5 × 5 min) and incubated in blocking solution (PBS, 0.1% bovine serum albumin, and 0.2% Triton X-100) for 1 hour. Next, tissue was placed in mouse anti-GAD67 overnight (described above), washed (5 × 5 min), and incubated in Cy3 donkey anti-mouse for 30 minutes. Tissue was then washed (5 × 5 min), slide mounted with polyvinyl medium, and coverslipped for imaging.

### 2.15 Microscopy

To determine injection placement, GFP was imaged with a Zeiss Axio Imager Z2 microscope using the 10x objective. To examine GFP co-localization with vGluT1, digital images were captured with optical sectioning (63x objective) to permit co-localization within a given z-plane (0.5-μm thickness). Co-localization was defined as yellow fluorescence from the overlap between labeled GFP terminals and red-colored vGluT1. The Zeiss Axio Imager Z2 microscope was also used to collect tiled brightfield images (10x objective) of ΔFosB-labeled tissue sections bilaterally. Imaging of dual-fluorescent ΔFosB and CaMKIIα or GAD67 was carried out with a 63x objective and optical sectioning (0.5-μm thickness) to determine if nuclear ΔFosB was surrounded by cytosolic markers for glutamatergic and/or GABAergic neurons within a 0.5-μm z-plane. SynaptoTag-injected tissue was imaged with the 10x objective to verify mCherry expression in the IL and GFP labeling in the insula. High-magnification images of GFP appositions onto CaMKIIα- or GAD67-labeled cell bodies in the insula were captured with a 63x objective using 0.5-μm thick optical sectioning.

### 2.16. Image analysis

Tiled-brightfield images were uniformly brightness- and contrast-corrected before anatomical delineation in ImageJ Fiji (ver. 1.51N). The bregma location of each tissue section was determined according to the Swanson atlas (Swanson, 2004) and images were hand delineated in ImageJ according to Swanson with supplementation from the Paxinos and Watson atlas (Paxinos & Watson, 2007) for divisions of the orbital cortex. Treatment-blinded observers delineated boundaries for the regions of interest, differentiating between superficial layers (1-3) and deep layers (4-6), as indicated by the Swanson atlas (Swanson, 2004). Regions analyzed were IL, prelimbic cortex (PL), anterior cingulate cortex (AC), medial orbital cortex (MO), ventral orbital cortex (VO), lateral orbital cortex (LO), anterior insular cortex (AI), and posterior insular cortex (PI). In the case of agranular cortex (e.g. IL), only layers 5 and 6 were included in the deep layers. For structures with distinct dorsal and ventral divisions (e.g. AI), these were analyzed separately. Following delineation, images were uniformly auto-thresholded for ΔFosB+ cells. Following thresholding, a watershed function and size-exclusion parameter were applied to specifically differentiate immuno-reactive nuclei. Cell counts for each region were then corrected for unit area.

### 2.17. Data analysis

Data are expressed as mean ± standard error of the mean. Data were analyzed using Prism 8 (GraphPad, San Diego, CA), with statistical significance set at p < 0.05 for all tests. The effects of CVS and vGluT1 siRNA treatment on the number of ΔFosB expressing cells were analyzed in the IL, PL, AC, MO, VO, LO, AI, and PI, using two-way analysis of variance (ANOVA) with distance from bregma (i.e. rostral-caudal location) and treatment as factors. In areas with a main effect of treatment, or an interaction between treatment and bregma location, Tukey’s multiple comparisons post-hoc analysis was conducted to determine the significant difference between groups at a given location. This analysis was also conducted within the superficial and deep layers of each region of interest, as well as dorsal and ventral areas of the AC and AI. Specific anatomical locations were only analyzed when samples were present from at least 3 animals in all groups. The mean number of ΔFosB-expressing nuclei did not differ between left and right hemispheres so bilateral measurements were pooled for analysis. Behavior in the FST was analyzed with an unpaired test comparing GFP and siRNA treatment within stress conditions. Behavior data throughout CVS were analyzed by two-way repeated-measures ANOVA with treatment and day (repeated) as factors. In the case of significant main effects, Sidak’s multiple comparisons post-hoc tests determined specific differences. For tone-shock conditioning, freezing was analyzed by two-way repeated-measures ANOVA with trial (repeated) and treatment as factors followed by Sidak’s multiple comparison post-hoc test. Data points significantly deviating from group mean (> 2 standard deviations) were uniformly identified with an Excel macro and excluded from analysis as outliers.

## 3. Results

### 3.1. Injection placement

Microinjections of the lentiviral-packaged construct expressing vGluT1 siRNA targeted the IL (Fig.1C&D). GFP expression was predominantly in the deep layers of the IL with minimal spread to the PL (Fig.1E). Injections with spread into more than 20% of the PL were excluded. The vGluT1 siRNA approach reduces vGluT1 mRNA in the IL, as well as vGluT1 protein on GFP-labeled terminals (Myers et al., 2017; Schaeuble et al., 2019). Here, the co-localization of immunolabeled vGluT1 with the GFP control construct was evident in cortico-cortical axonal processes (Fig.1F). However, vGluT1 protein was reduced on axonal processes in animals receiving the vGluT1 siRNA construct (Fig.1G).

### 3.2. Infralimbic cortex ΔFosB expression

A neuroanatomical survey of ΔFosB expression in the mPFC was conducted to localize the effects of chronic variable stress (CVS) and determine the role of decreased IL output. In the IL, treatment [F(2,82) = 5.841, p = 0.0043] and location effects [F(4,82)= 21.56, p < 0.0001] (Fig.2A) were found where CVS GFP rats (n = 9) had greater ΔFosB+ cell density than No CVS GFP rats (n = 7, p < 0.05) at 2.8 mm from bregma. Further, CVS siRNA rats (n = 9) had increased ΔFosB+ cell density compared to CVS GFP (p < 0.05, 3.2 mm) and No CVS GFP (p < 0.05, 2.8 and 2.15 mm). Superficial IL layers showed effects of group [F(2,80)= 13.86, p < 0.0001], location [F(4,80)= 81.69, p < 0.0001], and group × location interactions [F(8,80) = 5.493, p < 0.0001] (Fig.2B). Multiple comparisons indicated CVS GFP (n = 9) had increased ΔFosB+ cell density compared to No CVS GFP (n = 7, p < 0.05) at 2.8 mm from bregma. Additionally, CVS siRNA animals (n = 9) had higher ΔFosB+ cell density compared to CVS GFP (p < 0.05) at 3.2 mm and No CVS GFP in rostral portions (3.2 – 2.15 mm). In deep layers, there were treatment [F(2,80)= 7.353, p = 0.0012] and location effects [F(4,80) = 14.27, p < 0.0001] (Fig.2C). Specifically, CVS siRNA rats (n = 9) had higher ΔFosB+ density than CVS GFP (n = 9, p < 0.05, 3.2 mm) and No CVS GFP rats (n = 7, p < 0.05, 2.8 and 2.15 mm). Both generally and within superficial layers, CVS increased IL activation, an effect that was augmented by siRNA.

**Figure 2:**
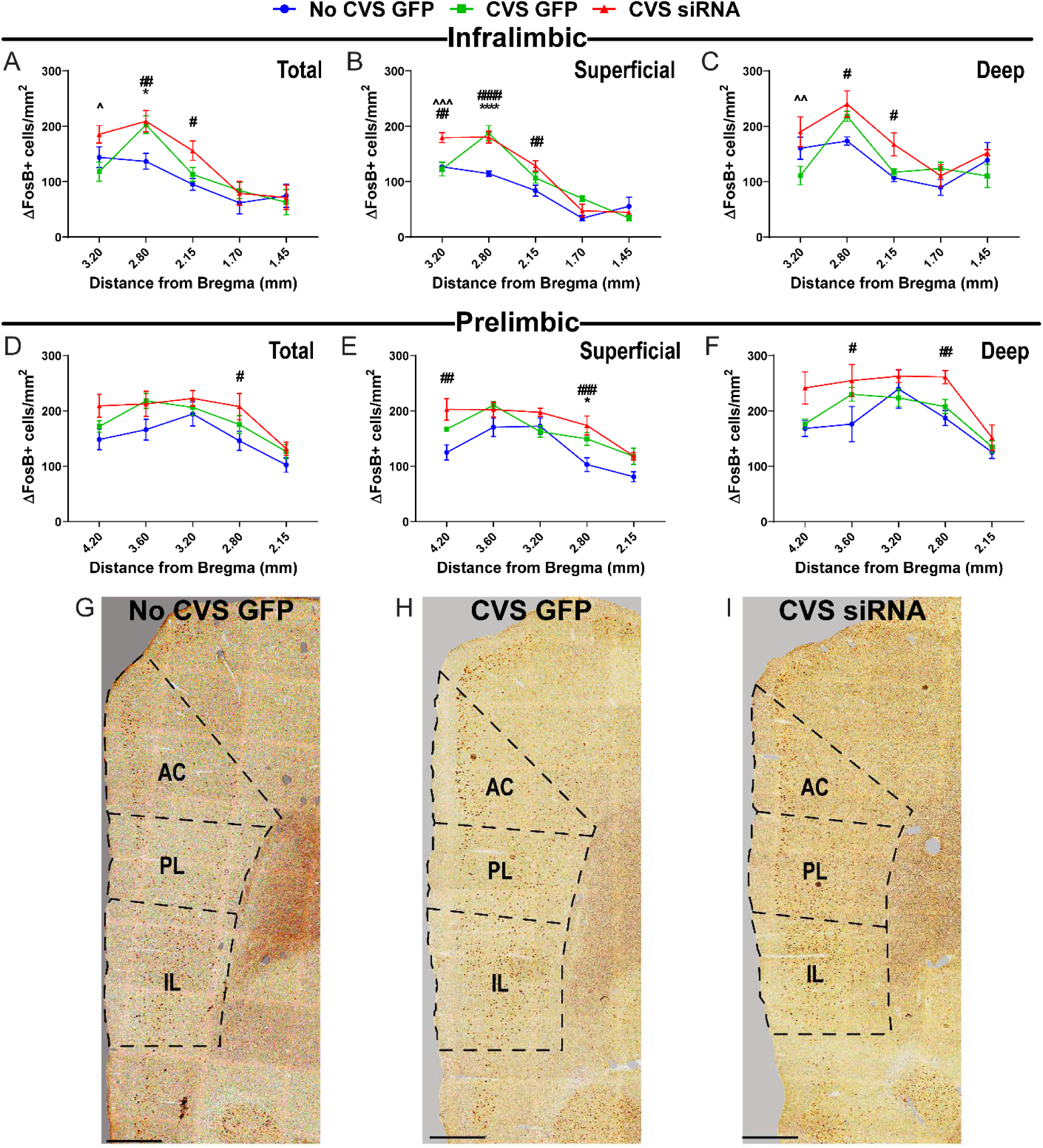
Limbic cortex ΔFosB expression. ΔFosB+ nuclei were increased by CVS GFP and CVS siRNA in the total IL (A), as well as superficial (B) and deep layers (C). ΔFosB-expressing cells were measured in the total PL (D), superficial (E), and deep layers (F). Representative images of the mPFC, including AC, PL, and IL at +2.8 mm from bregma from No CVS GFP (G), CVS GFP (H), and CVS siRNA (I) treatments, scale bars: 500 μm. Data are expressed as cell counts per mm^2^ ± SEM. * CVS GFP vs No CVS GFP, # CVS siRNA vs No CVS GFP, ^ CVS siRNA vs CVS GFP. *^#^^ p < 0.05, **^##^^^ p < 0.01, ***^###^^^^ p < 0.001, and ^^^^ p < 0.0001. IL: infralimbic cortex, PL: prelimbic cortex.

### 3.3. Prelimbic cortex

Spatial analysis of ΔFosB+ cell density in the PL (Fig.2D) indicated effects of group [F(2,82) = 8.303, p = 0.0005] and anatomical location [F(4,82) = 13.10, p < 0.0001]. Specifically, CVS siRNA animals (n = 9, p < 0.05) had greater ΔFosB+ density at +2.8 mm from bregma compared to No CVS GFP rats (n = 7, p < 0.05). In superficial PL layers, group [F(2,78) = 18.81, p < 0.0001] and location effects [F(4,78)= 26.22, p < 0.0001] were found (Fig.2E) indicating CVS (n = 9, p < 0.05) increased ΔFosB compared to No CVS GFP rats (n = 7) at 2.8 mm from bregma. Also, CVS siRNA (n = 9, p < 0.05) increased expression across superficial (4.2 and 2.8) and deep (3.6 and 2.8) layers where effects of group [F(2,80) = 11.11, p < 0.0001] and location [F(4,80) = 15.73, p < 0.0001] (Fig.2F) were found. While CVS increased activation in superficial PL, more widespread PL increases were seen with CVS siRNA.

### 3.4 Anterior cingulate cortex

Across the total AC there were main effects of treatment [F(2,205) = 5.13, p = 0.0067] and location [F(14,205) = 46.39, p < 0.0001] (Fig.3A). Post-hoc analysis indicated that CVS siRNA rats (n = 9) had an increase in ΔFosB+ nuclei compared to No CVS GFP (n = 7, p < 0.05) and CVS GFP (n = 9, p < 0.05) rats in the rostral portion of the AC (3.6 – 2.15 mm from bregma). Within the dorsal division of the AC, there were main effects of treatment [F(2,203) = 6.11, p = 0.0026] and location [F(14,203) = 33.13, p < 0.0001], as well as a treatment × location interaction [F(28,203) = 1.63, p = 0.0292] (Fig.3B). Specifically, CVS siRNA rats (n = 9) had increased ΔFosB+ cell density compared to No CVS GFP (n = 7, p < 0.05) and CVS GFP (n = 9, p < 0.05) in rostral AC regions (3.2 – 2.15 mm from bregma). In the ventral AC, 2-way ANOVA found an effect of location [F(9,122)= 29.19, p < 0.0001] but no treatment effect (Fig.3C).

**Figure 3:**
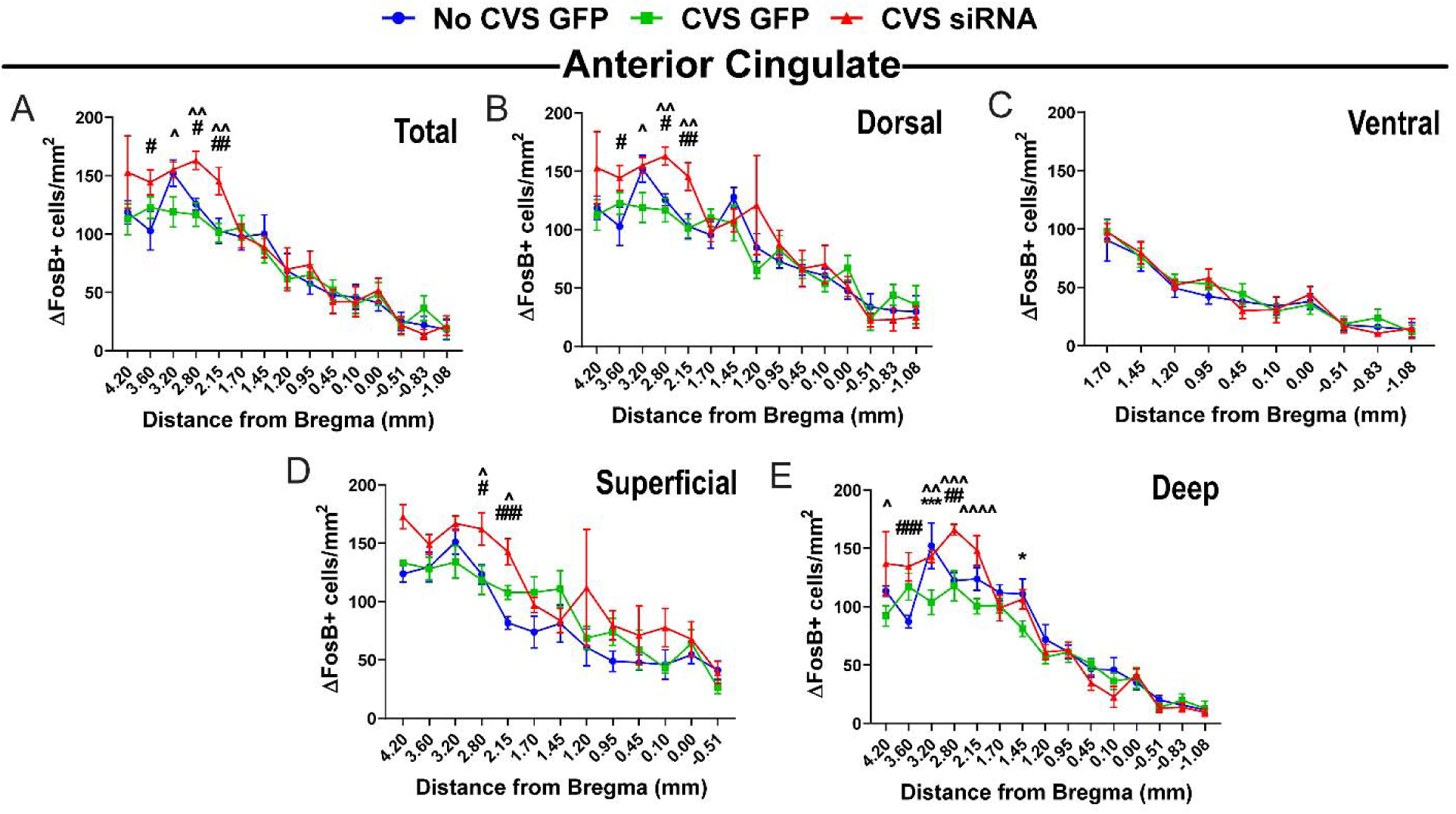
Cingulate ΔFosB expression. ΔFosB+ cell density quantified throughout the total AC (A), the dorsal division (B), and the ventral portion (C). Immunoreactive cells were also quantified in the superficial (D) and deep layers (E). The CVS siRNA group had increased ΔFosB density compared to No CVS GFP and CVS GFP in the rostral portions of the AC (n = 7­9/group). Data are expressed as cell counts per mm^2^ ± SEM. * CVS GFP vs No CVS GFP, # CVS siRNA vs No CVS GFP, ^ CVS siRNA vs CVS GFP. *^#^^ p < 0.05, **^##^^^ p < 0.01, ***^###^ p < 0.001, and ****^####^ p < 0.0001. AC: anterior cingulate cortex.

Density of ΔFosB expression was also determined in superficial (I-III) and deep (IV-VI) layers. Superficial layers had a treatment [F(2,181) = 15.61, p < 0.0001] and location effect [F(12,181) = 26.77, p < 0.0001] (Fig.3D) localized by post-hoc analysis to rostral AC (2.8 – 2.15 mm to bregma) where CVS siRNA animals (n = 9) had higher ΔFosB+ cell density than No CVS GFP (n = 7, p < 0.05) and CVS GFP (n = 9, p < 0.05). In the AC deep layers, there was a location effect [F(14,203) = 67.84, p < 0.0001] and treatment × location interaction [F(28,203) = 2.580, p < 0.0001] (Fig.3E). Here, CVS decreased ΔFosB+ cell density compared to no CVS GFP (p < 0.05) at 3.2 and 1.45 mm from bregma. However, CVS siRNA rats (n = 9) had increased cell density compared to both CVS GFP (n = 9, p < 0.05) and no CVS GFP rats (n = 7, p < 0.05). Other than CVS effects to decrease chronic activation in deep layers of AC, the general effect of this quantification was increased AC activation in CVS siRNA animals compared to the other groups.

### 3.5. Medial orbital cortex

In the orbital cortices, 2-way ANOVA of total MO ΔFosB+ nuclei did not find significant effects (Fig.4A). However, analysis of MO superficial layers revealed a treatment effect [F(2,24)=4.352, p=0.0244] (Fig.4B). Post-hoc comparisons found CVS siRNA rats (n = 9) had increased ΔFosB+ cell density compared to No CVS GFP (n = 7, p < 0.05) at 4.2 mm rostral to bregma. Analysis of deep layers indicated an anatomical location effect [F(1,25)=5.126, p=0.0325] (Fig.4C); however, there were no specific location or treatment effects in multiple comparisons. Overall, the only MO changes were in the superficial layers of CVS siRNA animals.

**Figure 4:**
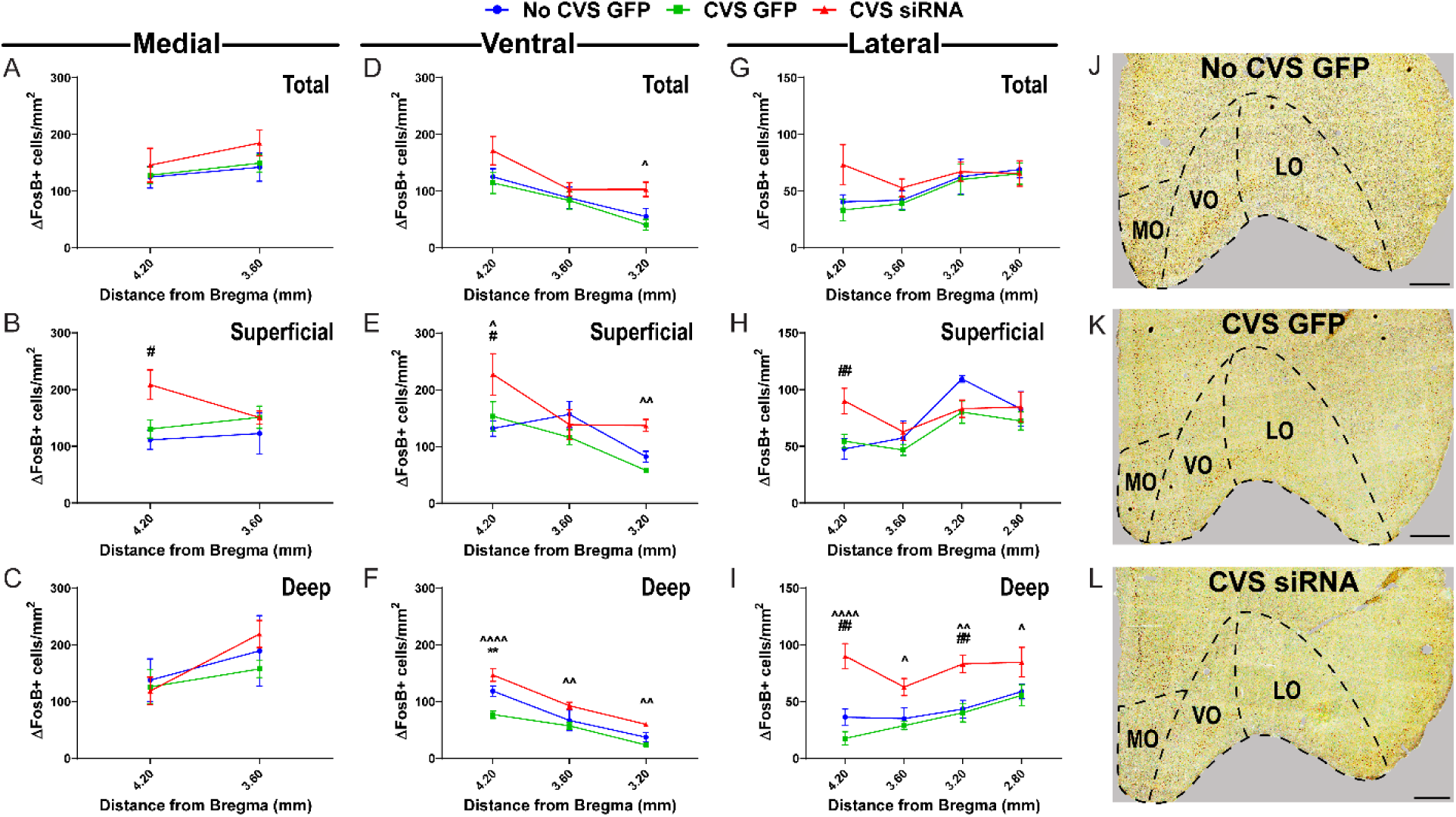
Orbital cortex ΔFosB expression. ΔFosB+ cell density measured through the total MO (A), along with the superficial (B) and deep layers (C). The density of ΔFosB immunoreactivity was surveyed throughout the total (D), superficial (E), and deep layers of the VO (F). ΔFosB+ cell density was also quantified throughout the total (G), superficial (H), and deep layers of the LO (I). ΔFosB+ cell densities were increased in CVS siRNA rats compared to CVS GFP (n = 7-9/group) throughout multiple orbital regions. Representative images of the orbital cortices, including MO, VO, and LO at +3.6 mm from bregma from No CVS GFP (J), CVS GFP (K), and CVS siRNA (L) treatments, scale bar: 500 μm. Data are expressed as cell counts per mm^2^ ± SEM. * CVS GFP vs No CVS GFP, # CVS siRNA vs No CVS GFP, ^ CVS siRNA vs CVS GFP. *^#^^ p < 0.05, **^##^^^ p < 0.01, ^^^ p < 0.001, and ^^^^ p < 0.0001. MO: medial orbital cortex, VO: ventral orbital cortex, LO: lateral orbital cortex.

### 3.6. Ventral orbital cortex

Analysis of ΔFosB-expressing cells/mm^2^ in the VO found an effect of treatment [F(2,42) = 6.949, p = 0.0025] and location [F(2,42) = 13.80, p < 0.0001] (Fig.4D). Specifically, CVS siRNA rats (n = 9) had increased ΔFosB+ cell density compared to No CVS GFP (n = 7, p < 0.05) at 3.2 mm from bregma. Analysis of superficial VO regions showed treatment [F(2,40) = 7.737, p = 0.0014] and location effects [F(2,40) = 13.01, p < 0.0001] (Fig.4E). Here, CVS siRNA rats (n = 9) had higher ΔFosB+ cell density than No CVS GFP (n = 7, p < 0.05, 4.2 mm) and CVS GFP (n = 9, p < 0.05, 4.2 and 3.2 mm). Analysis of deep VO populations indicated group [F(2,41) = 23.33, p < 0.0001] and location effects [F(2,41) = 52.01, p < 0.0001] (Fig.4F). Here, CVS decreased activation in the rostral portion while CVS siRNA rats (n = 9) had a higher cell density than CVS GFP (n = 9, p < 0.05) throughout the deep VO. Collectively, CVS decreased activation specifically in deep layers of VO, while CVS and siRNA treatment led to increases throughout.

### 3.7. Lateral orbital cortex

Analysis of the total LO showed anatomical location effects [F(3,62) = 3.481, p = 0.021] (Fig.4G) without multiple comparison significance. However, LO superficial layers showed an effect of location [F(3,60)= 6.630, p=0.0006] (Fig.4H) and multiple comparisons revealed CVS siRNA (n = 9) had increased ΔFosB+ density compared to No CVS GFP (n = 7, p < 0.05) rostrally (4.2 mm). Furthermore, deep-layered LO had treatment [F(2,60) = 26.93, p < 0.0001] and location effects [F(3,60) = 4.490, p = 0.0066] (Fig.4I) where CVS siRNA rats (n = 9) had increased ΔFosB+ cell density across all rostral-caudal locations. Thus, CVS siRNA increased LO activation.

### 3.8. Anterior insular cortex

Analysis of the AI by 2-way ANOVA showed treatment [F(8,139)=11.01, p<0.0001] and location [F(2,139)=14.44, p<0.0001] effects (Fig.5A). Here, there were decreased ΔFosB+ cells in CVS GFP rats (n = 9) and increased cell density in CVS rats treated with siRNA (p < 0.05). For dorsal AI, treatment [F(2,139) = 19.70, p < 0.0001] and location effects [F(8,139) = 17.09, p < 0.0001] (Fig.5B) were present where CVS GFP rats (n = 9) exhibited decreased ΔFosB+ cells, an effect prevented by siRNA (p < 0.05). Similarly, ventral AI showed treatment [F(2,96) = 13.05, p < 0.0001] and location effects [F(5,96) = 5.219, p = 0.0003] (Fig.5C) with CVS GFP rats (n = 9) again exhibiting lower ΔFosB+ counts compared to controls. In contrast, CVS siRNA (n = 9, p < 0.05) increased activation compared to CVS GFP. Layer-specific analyses revealed treatment and location effects in the superficial (Fig.5D) ([F(2,139) = 39.09, p < 0.0001] and [F(8,139)=5.737, p<0.0001], respectively) and deep (Fig.5E) ([F(2,138) = 12.31, p < 0.0001] and [F(8,138) = 20.73, p < 0.0001], respectively) AI. In both cell groups CVS GFP animals (n = 9) had lower ΔFosB+ densities compared to No CVS GFP (n = 7, p < 0.05), while CVS siRNA (n = 9, p < 0.05) had more expression across the rostral-caudal gradient compared to both No CVS GFP and CVS GFP. Generally, CVS decreased AI activation, an effect that was prevented by siRNA treatment particularly in the superficial layers.

**Figure 5:**
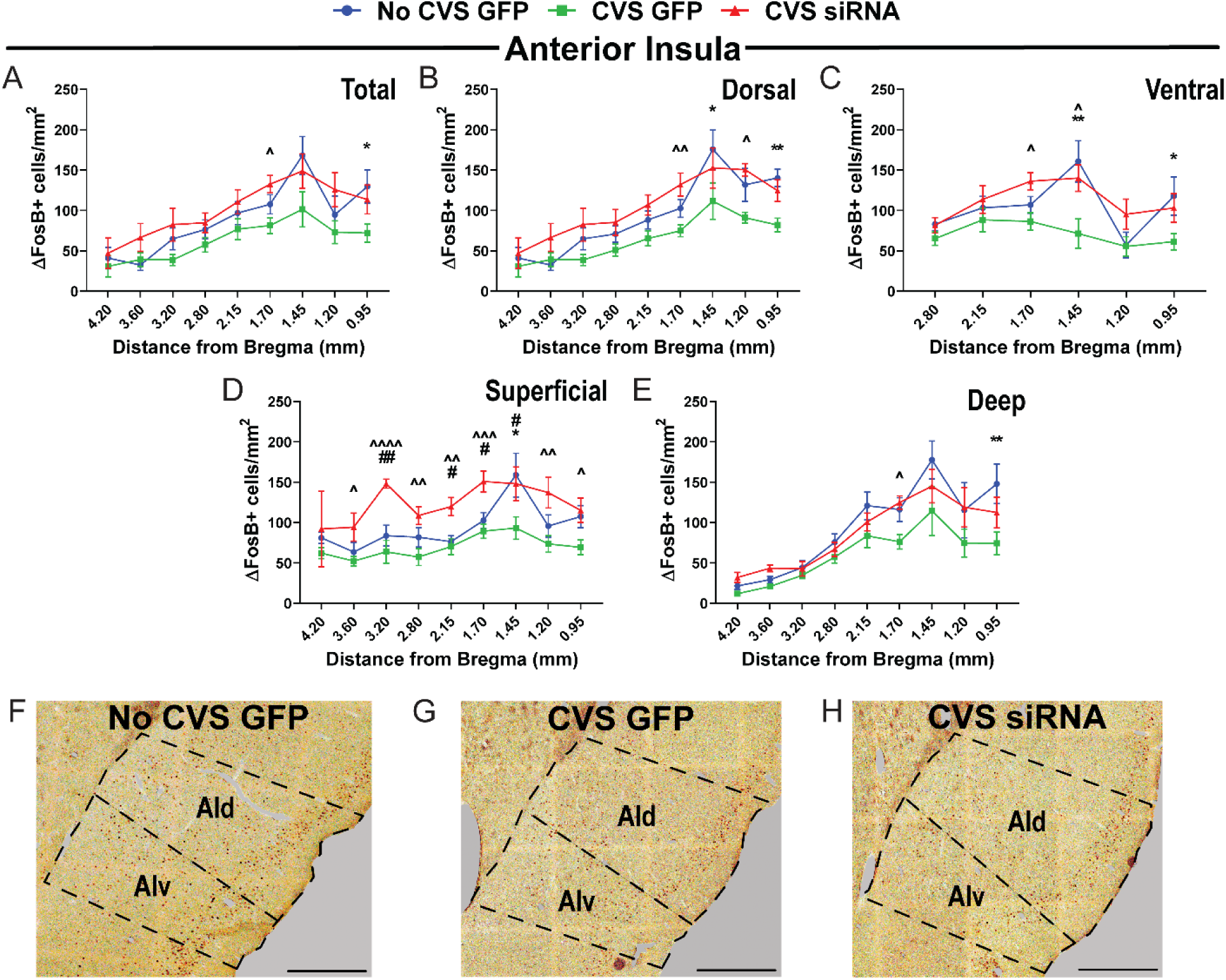
Anterior insula ΔFosB expression. ΔFosB immunoreactivity was quantified in total AI (A), dorsal region of the AI (B), ventral region of the AI (C), as well as superficial (D) and deep layers (E) of the AI. Generally, cell counts were decreased in CVS GFP, an effect prevent by CVS siRNA (n = 7-9/group). Representative images of the AI at +1.4 mm from bregma from No CVS GFP (F), CVS GFP (G), and CVS siRNA (H) treatments, scale bar: 500 μm. Data are expressed as cell counts per mm^2^ ± SEM. * CVS GFP vs No CVS GFP, # CVS siRNA vs No CVS GFP, ^ CVS siRNA vs CVS GFP. *^#^^ p < 0.05, **^##^^^ p < 0.01, ^^^ p < 0.001, and ^^^^ p < 0.0001. AId: anterior insular cortex, dorsal part, AIv: anterior insular cortex, ventral part.

### 3.9. Posterior insular cortex

Across the PI effects of treatment [F(2,186) = 18.13, p < 0.0001] and location [F(12,186) = 16.44, p < 0.0001] were present (Fig. 6A). Post-hoc comparisons showed CVS GFP animals (n = 9) had decreased ΔFosB+ density compared to No CVS GFP (n = 7, p < 0.05) at numerous locations and CVS siRNA (n = 9, p < 0.05) had an increase in ΔFosB-expressing cells. This trend continued in superficial and deep cells of the PI. Specifically, superficial PI had effects of group [F(2,182) = 20.76, p < 0.0001] and location [F(12,182) = 4.52, p < 0.0001] (Fig. 6B) as did PI deep layers (group: [F(2,186) = 11.48, p < 0.0001], location: [F(12,186) = 15.90, p < 0.0001]) (Fig. 6C). Post-hoc analysis further indicated that, at multiple locations, CVS GFP animals had lower ΔFosB+ cell density while CVS siRNA animals had increased ΔFosB immunoreactivity. Similar to AI, PI showed CVS-induced decreases in activation that were IL-dependent.

**Figure 6:**
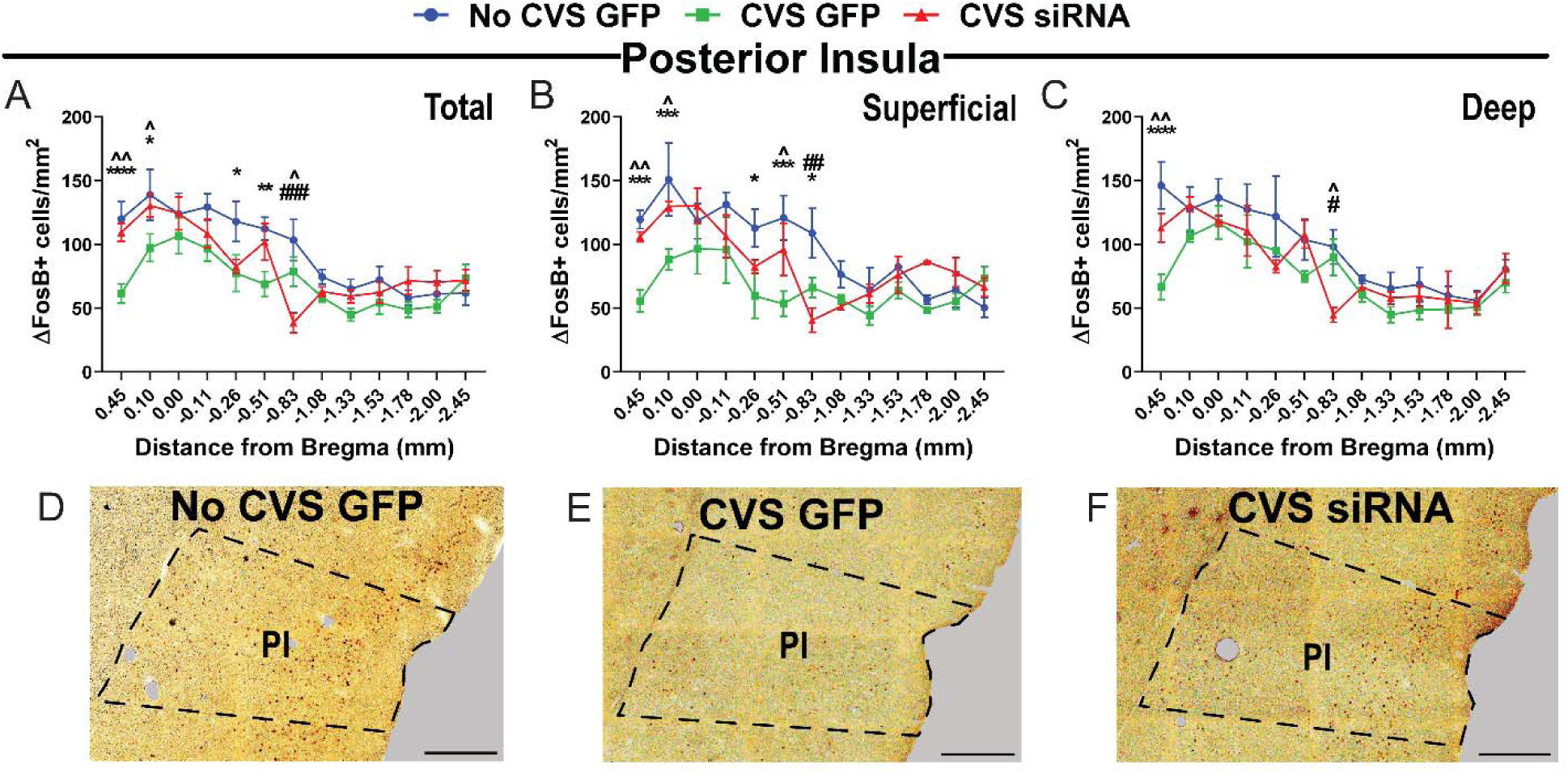
Posterior insula ΔFosB expression. ΔFosB+ cell density was measured throughout the total PI (A), along with superficial (B) and deep layers (C). CVS GFP groups had less ΔFosB+ cell density compared to No CVS GFP (n = 7-9/group) and CVS siRNA (n = 7-9/group). Representative images of the AI at +1.4 mm from bregma from No CVS GFP (D), CVS GFP (E), and CVS siRNA (F) treatments, scale bar: 500 μm. Data are expressed as cell counts per mm^2^ ± SEM. * CVS GFP vs No CVS GFP, # CVS siRNA vs No CVS GFP, ^ CVS siRNA vs CVS GFP. *^#^^ p < 0.05, **^##^^^ p < 0.01, ***^###^ p < 0.001, and **** p < 0.0001. PI: posterior insular cortex.

### 3.10. Neurochemical phenotype of ΔFosB-expressing cells

Given the CVS effects on activity patterns in IL and AI, we sought to determine whether these changes were localized to excitatory and/or inhibitory cells. Accordingly, ΔFosB co-expression with either CaMKIIα or GAD67 was examined in prefrontal and insular tissue of CVS-exposed animals. Co-localization with the pyramidal neuron marker CaMKIIα indicated that at most ΔFosB expression was in excitatory neurons (Fig.7A&C). In contrast, examination of interneuronal GAD67+ cells found a lack of co-localization with ΔFosB throughout prefrontal and d insular cortices (Fig.7B&D).

**Figure 7:**
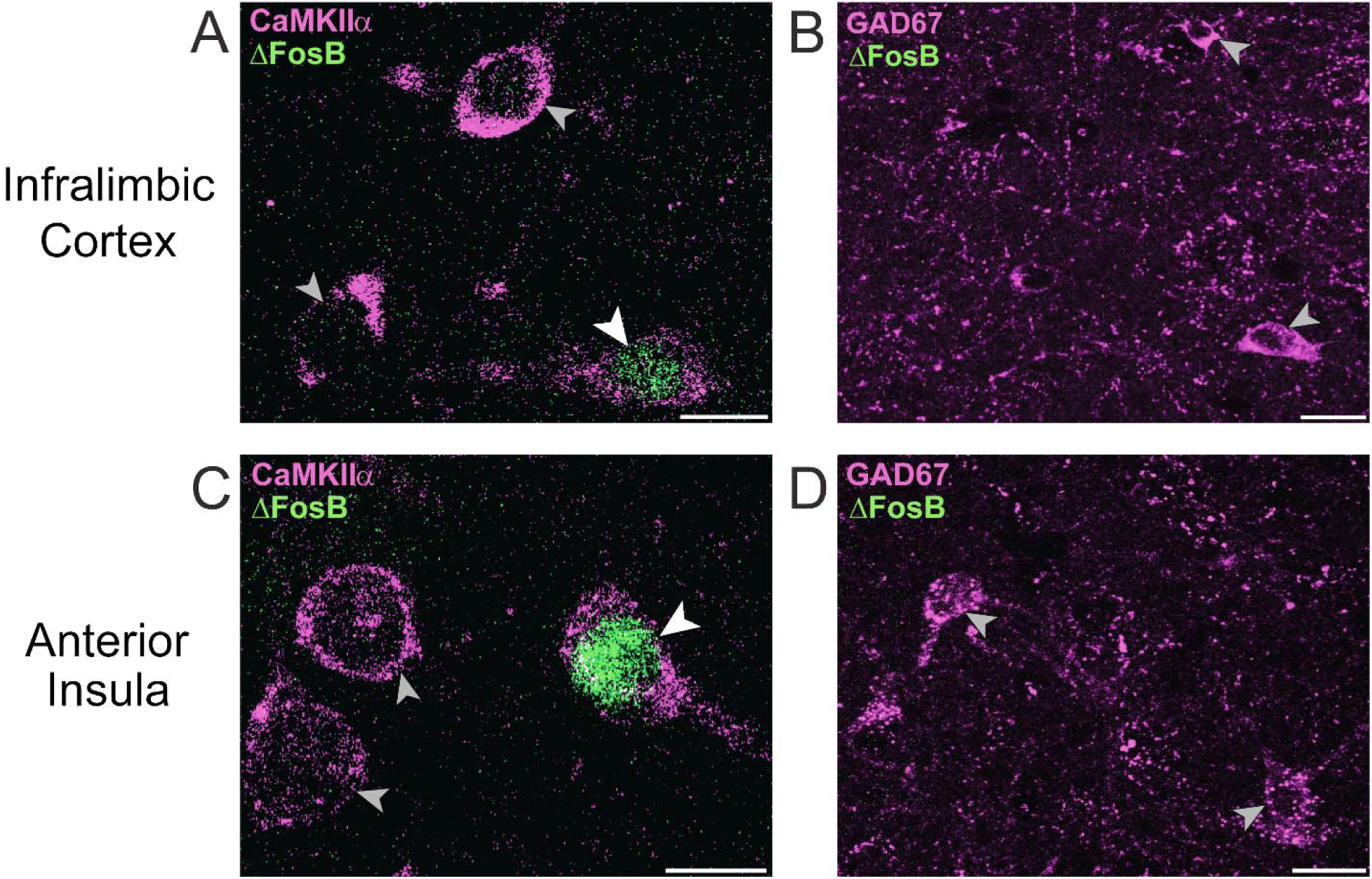
ΔFosB is expressed in pyramidal neurons but not interneurons in the IL and AI. Dual immunofluorescence in layer III of the IL and AI identified ΔFosB cells co-labeled with CaMKIIα (A, C), scale bar: 10 μm. ΔFosB did not label GAD67-positive neurons in either the IL or AI (B, D), scale bar: 10 μm. White arrowheads: representative co-labeled cells, grey arrowheads: representative non-ΔFosB+ neurons. CaMKIIα: Ca2+/calmodulin-dependent protein kinase II alpha, GAD67: glutamic acid decarboxylase, 67 kDa isoform, IL: infralimbic cortex, AI: anterior insular cortex.

### 3.11. Behavior

To investigate whether IL output influences the behavioral consequences of chronic stress, we examined multiple behaviors in animals undergoing CVS in experiments 1 and 2. Chronic stress regimens (CVS, chronic unpredictable stress, and chronic mild stress) are commonly used in rodents to model anxiety-like avoidance behaviors (Kaufmann & Brennan, 2018; Lezak et al., 2017; Sequeira-Cordero et al., 2019) and depression-like passive coping (Lam et al., 2018; Molina et al., 1994; Sequeira-Cordero et al., 2019; Willner, 2017). In the current studies, we assessed the contribution of IL glutamate to chronic stress-induced avoidance and coping behaviors.

### 3.12. Forced swim test

Studies were designed so that previously unstressed rats and CVS-exposed rats were subject to the FST in separate experiments. Prior to CVS, siRNA rats (n = 12) had decreased immobility [t(20) = 2.8, p = 0.012] (Fig.8A) and increased active coping (swimming and climbing) relative to GFP controls (n = 10) [t(20) = 2.6, p = 0.018] (Fig.8B). Similarly, in CVS rats, siRNA decreased immobility (n = 11) [t(23) = 2.2, p = 0.039] (Fig.8C) and increased activity [t(22) = 2.3, p = 0.034)] (Fig.8D). Collectively, these data indicate that attenuated IL glutamate output increases active coping regardless of stress history.

**Figure 8:**
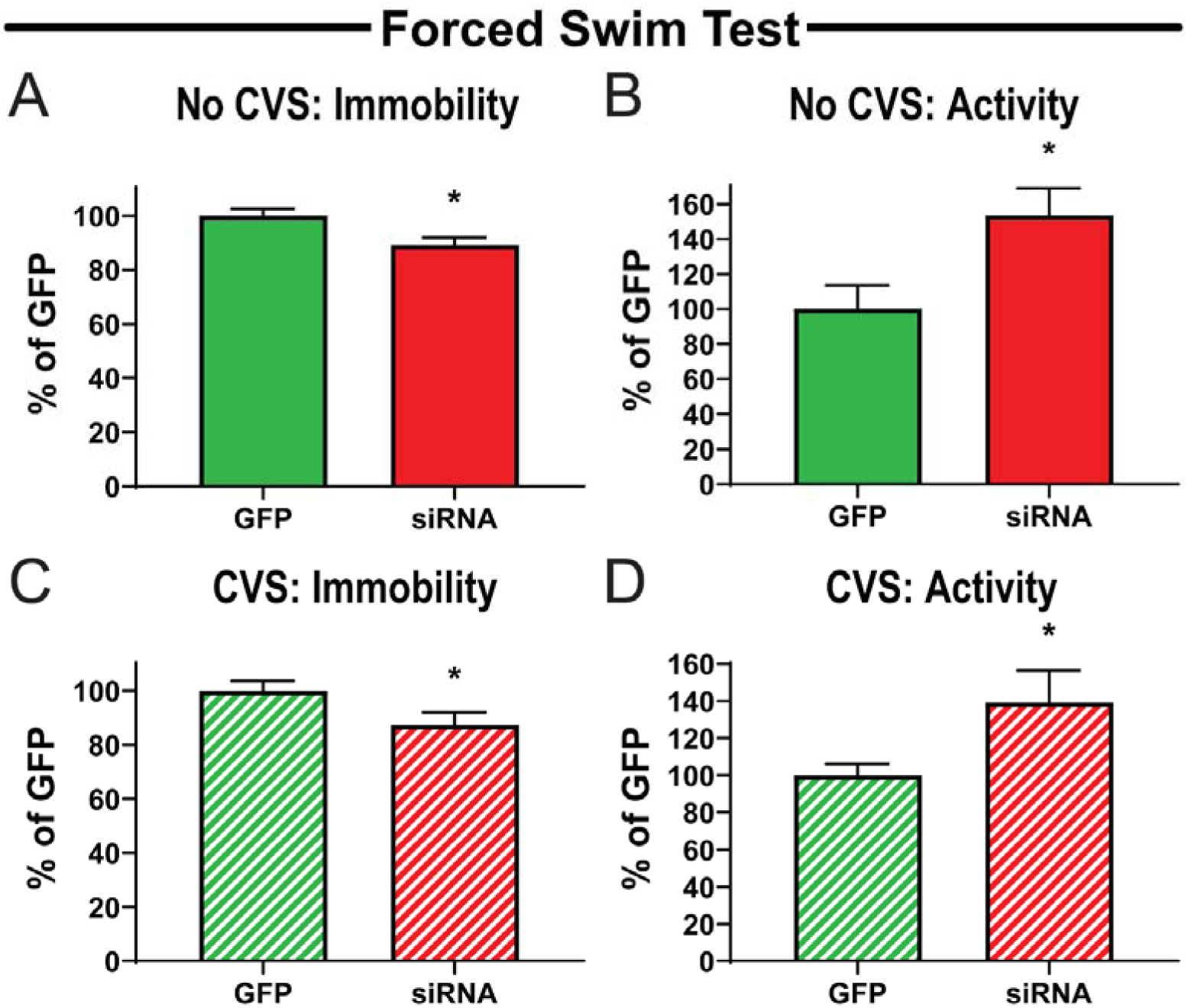
vGluT1 siRNA treatment reduces passive coping in previously unstressed and chronically stressed rats. No CVS siRNA rats had reduced immobility (A) and increased activity (B) in the forced swim test compared to No CVS GFP (n = 10-12). CVS siRNA animals also exhibited decreased immobility (C) and enhanced activity (D) compared to CVS GFP (n = 11-14). Data are expressed as percentage of GFP control ± SEM. * p < 0.05 compared to GFP.

### 3.13. Open field test/novel object interaction

Time spent in the center of the open field exhibited a main effect of day [F(1,22) = 8.76, p = 0.007] (Fig.9A). Post-hoc analysis found that CVS GFP rats (n = 13) spent less time (p = 0.025) in the center of the open field on day 13 compared to day 1. Notably, these effects were not present in siRNA-treated animals (n = 11). There were no significant effects on total distance traveled in the open field, indicating no CVS or siRNA effects on general locomotion (Fig.9B). Rats were also allowed to interact with a novel object in the center of the open field.

**Figure 9:**
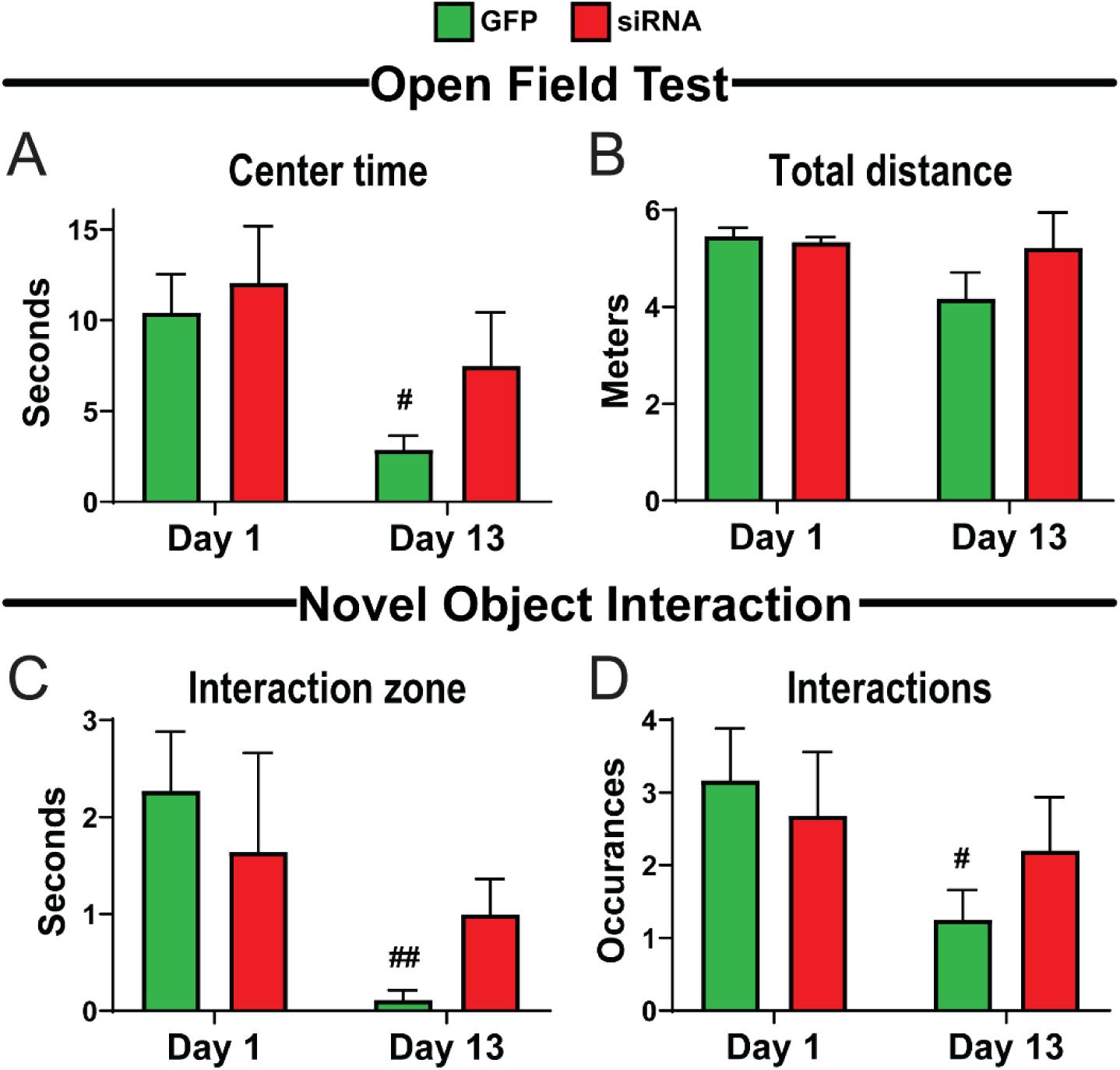
vGluT1 siRNA treatment prevents CVS effects on exploratory behavior. In the open field test, GFP rodents 13 days into CVS had reduced center time (A) (n = 11-14/group). No locomotion differences were observed in any condition (B). Time spent in the novel object interaction zone (C) and number of interactions (D) was reduced in GFP rats during CVS. Data are expressed as counts ± SEM. ^#^ p < 0.05, ^##^ p < 0.01 compared to Day 1.

Here, CVS affected time spent in the interaction zone [F(1,22) = 7.57, p = 0.012] (Fig.9C) and the frequency of object interactions [F(1,22) = 5.12, p = 0.034] (Fig.9D). Specifically, CVS GFP rats (n = 13) spent less time in the interaction zone (p = 0.009) and had fewer interactions (p = 0.027) on day 13 compared to day 1. However, siRNA rats (n = 11) did not have any significant behavioral change during CVS. Taken together, these results suggest that decreased IL glutamate output prevents stress-induced effects on approach-avoidance behaviors.

### 3.14. Elevated plus maze

Time spent on the open arms of the elevated plus maze was not altered by CVS or siRNA (Fig.10A). Although, the number explorations of the ends of the open arms had a treatment effect [F(1,20) = 10.30, p = 0.004] (Fig.10B). There was a significant increase in open arm end explorations for siRNA-treated rats (n = 11) compared to GFP controls (n = 8-9) on both day 1 (p = 0.011) and day 14 (p = 0.041). This effect of siRNA treatment may point to behavioral disinhibition and/or reduced risk aversion, regardless of stress.

**Figure 10:**
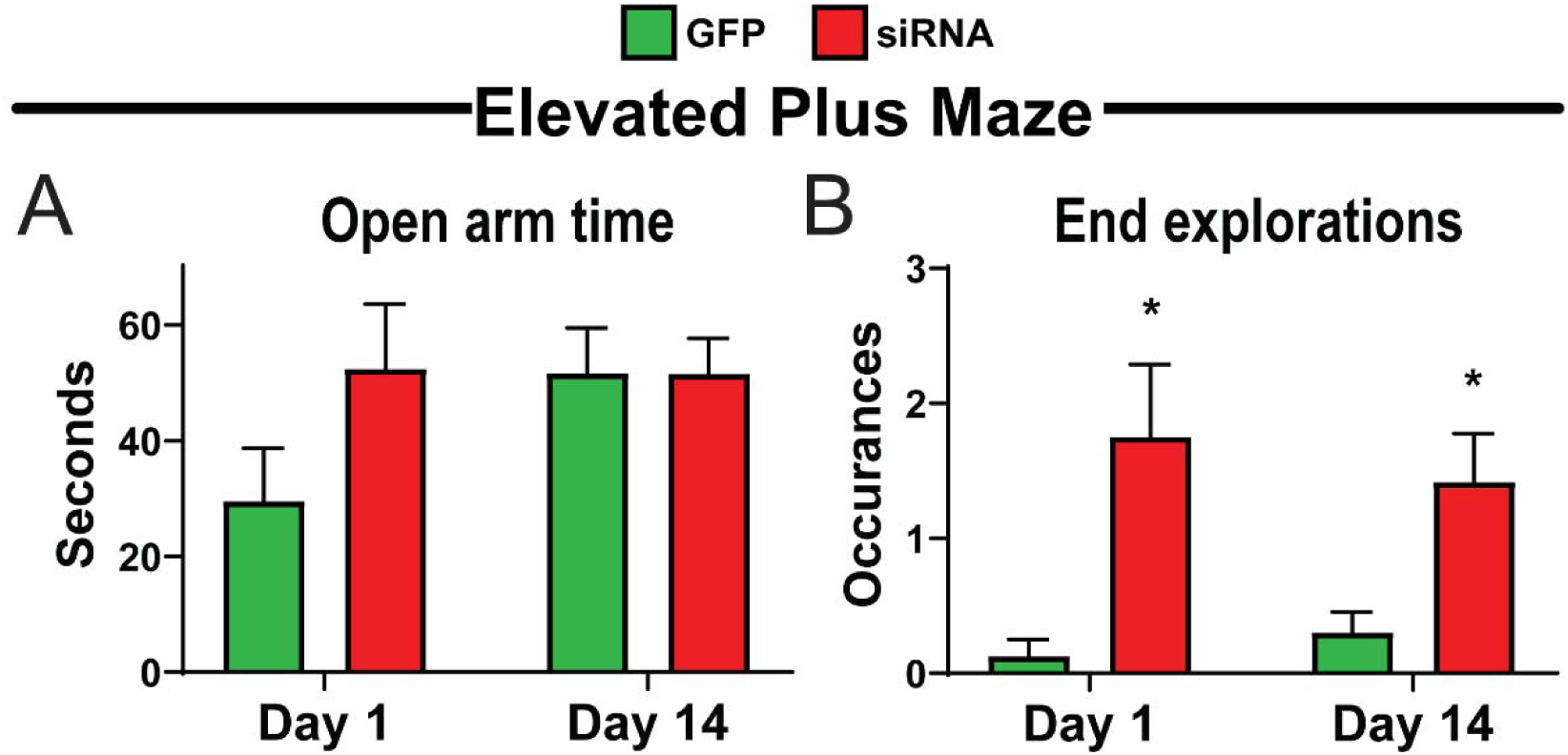
vGluT1 siRNA treatment increases end explorations of the elevated-plus maze. There were no effects of time in the open arm of the elevated-plus maze (A) (n = 8-12/group). siRNA rats explored the outer end of the open arm more on both days 1 and 14 (B). Data are expressed as counts ± SEM. * p < 0.05 compared to GFP.

### 3.15. Shock probe defensive burying

In response to the presence of the shock probe, there were significant day [F(1,25) = 22.18, p < 0.0001] and treatment [F(1,25) = 18.44, p = 0002] effects on probe contacts (Fig.11A). Post-testing revealed that siRNA treatment (n = 14) increased contacts on day 2 (p = 0.018). Further, on day 12 of CVS both treatment groups (n = 12-13) had reduced probe contacts (p < 0.01). However, there was still an effect of treatment where the siRNA had increased probe contacts (p=0.027). After the first probe contact, there were day [F(1,21) = 10.26, p = 0.004], treatment [F(1,25) = 11.18, p = 0.003], and day × treatment [F(1,21) = 4.47, p = 0.047] effects on the occurrences of stretch-attend behaviors toward the probe (Fig.11B). On day 12, siRNA animals exhibited stretch-attend posture significantly more than all other groups CVS (p < 0.01). In terms of defensive freezing after the first probe contact, both day [F(1,46) = 19.05, p < 0.0001] and treatment [F(1,46) = 8.81, p = 0.005] had significant effects (Fig.11C).

**Figure 11:**
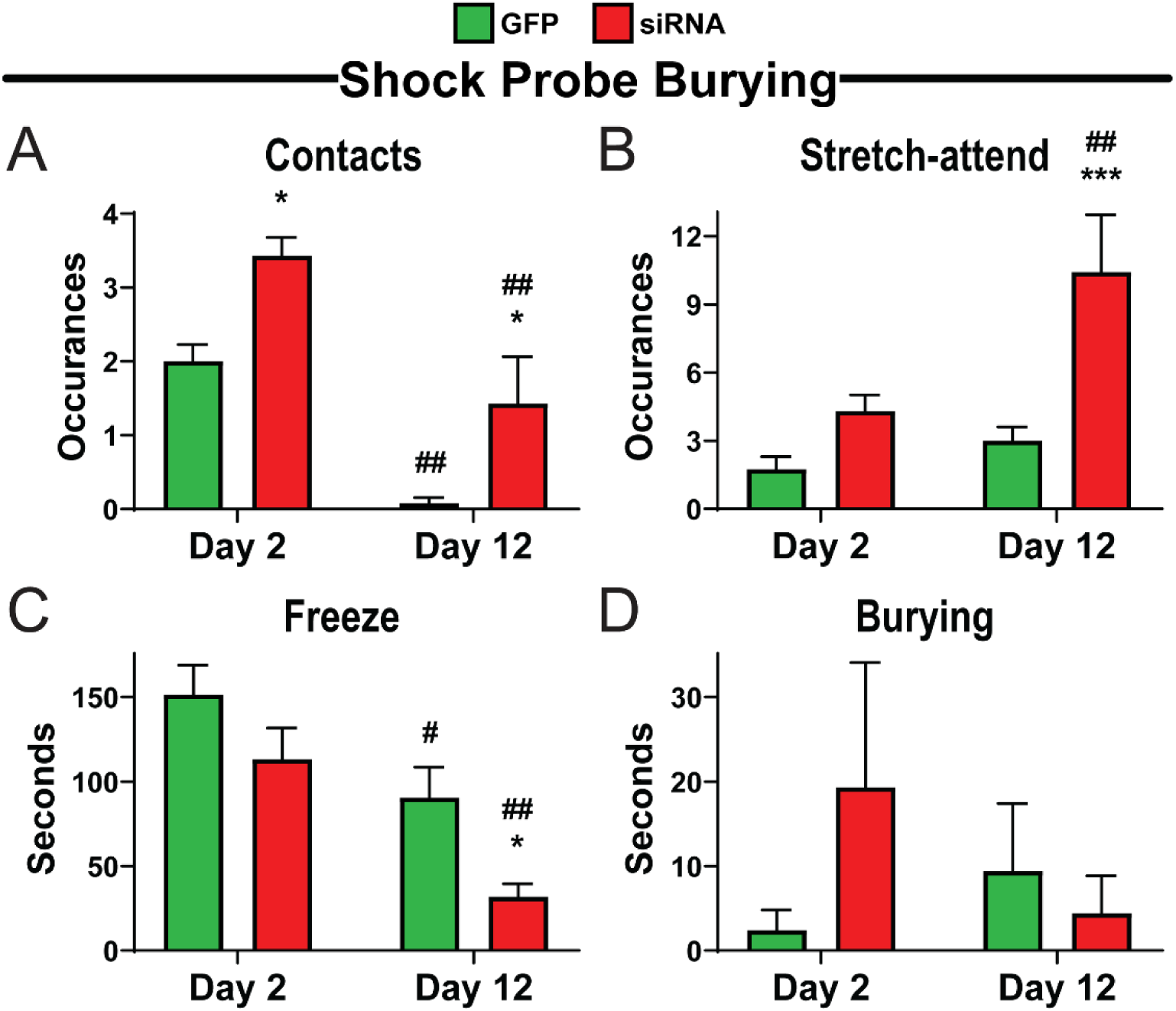
vGluT1 siRNA treatment increases shock probe interactions. siRNA-treated animals contacted the shock probe more times than GFP controls (A) (n = 8-12/group). On day 12, siRNA rats displayed more stretch-attend behaviors toward the probe (B). Freezing behavior decreased in both groups after 12 days of CVS, with siRNA rats freezing less (C). No significant changes were observed in shock probe burying (D). Data are expressed as counts ± SEM. * p < 0.05, *** p < 0.001 compared to GFP. ^#^ p < 0.05, ^##^ p < 0.01 compared to day 2.

On CVS day 12, both treatments spent less time freezing than day 2 (p < 0.05). Additionally, on day 12 siRNA animals froze less than GFP animals (p = 0.028). There were no significant effects of day or treatment on active burying behaviors (Fig.11D). Overall, vGluT1 siRNA reduced threat aversion and prevented avoidant and passive coping in a stress-dependent manner.

### 3.16. Von Frey assay

Given the lack of aversion that siRNA rats had for the shock probe, we assessed whether there were effects on nociception. However, there were no significant effects of day or treatment (n = 8-12/group) on somatic sensitivity (Fig.12A), suggesting that altered responses to aversive stimuli were not dependent on changes in nociception or pain sensitivity.

**Figure 12:**
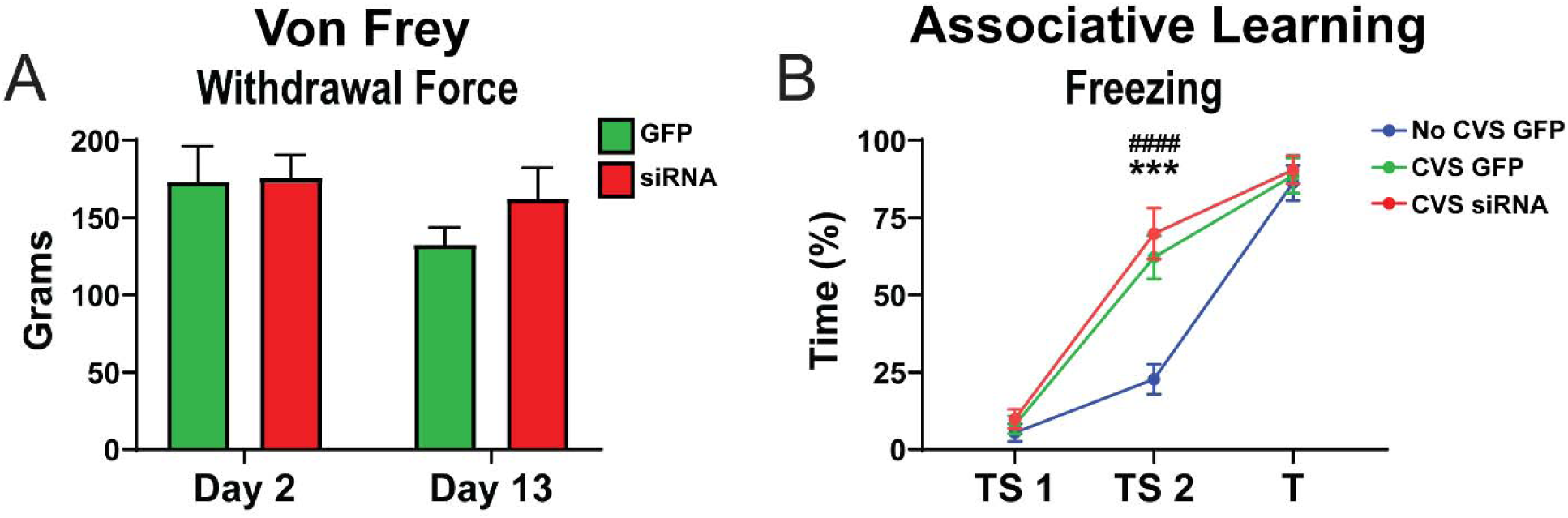
vGluT1 siRNA treatment does not impair nociception or associative learning. The Von Frey assay of somatic nociception found no changes in pain perception (A). Tone-shock conditioning found increased freezing in CVS GFP and CVS siRNA rats (n = 8-12f/group) during the second tone but all groups showed learned fear responses by the third tone (B). Data are expressed as counts and percentages ± SEM. *** p < 0.001 CVS GFP vs. No CVS GFP, ^####^ p < 0.0001 CVS siRNA vs. No CVS GFP. T: tone only, TS: tone-shock pairing.

### 3.17. Associative learning

Based on the effects of vGluT1 knockdown to reduce behavioral change during CVS and decrease aversion for noxious stimuli, we sought to determine how the siRNA may affect associative learning during CVS. An abbreviated fear conditioning protocol was used to examine IL effects on aversive associations. Time spent freezing during tone presentation was measured preceding 2 shocks and for a third tone that was not shock-paired. Repeated measures ANOVA found effects of trial [F(2,68) = 184.7, p < 0.0001], treatment [F(3,34) = 4.357, p = 0.0106], and a trial × treatment interaction [F(6,68) = 3.230, p = 0.0075] (Fig.12B). Post-hoc comparisons found that, during the second tone, CVS-exposed animals (n = 10) had more freezing than No CVS GFP (n = 8) animals regardless of treatment (p < 0.01). Ultimately, all groups demonstrated equivalent fear learning by the third trial. These data suggest that CVS siRNA rats (n = 12) don’t have deficits in associative conditioning that could account for the reduced behavioral change during CVS.

### 3.18. Infralimbic cortex-anterior insula circuit

Given our finding that CVS decreased ΔFosB expression in the insula in an IL-dependent manner, we sought to examine direct IL synaptic projections to the AI. Further, we determined the post-synaptic chemistry of IL-targeted insular cells to better understand potential effects on excitatory/inhibitory balance. The SynaptoTag virus was injected into the IL which leads to mCherry expression in cell bodies and axons of transduced neurons (n = 4) (Fig.13A). Further, conjugation of GFP to synaptobrevin-2 leads to GFP expression specifically at pre-synaptic terminals. This approach allows examination of IL efferents forming putative synapses without concern for fibers-of-passage. Microinjection placement (Fig.13B), evidenced by somatic mCherry, was confined to the IL (Fig.13C). Low-magnification micrographs in the AI indicated GFP-labeled IL synapses in both superficial and deep layers (Fig.13D). Moreover, AI neurons immuno-labeled for CaMKIIα and GAD67 in layer VI were apposed by GFP-labeled IL synaptic terminals (Fig.13E&F). Taken together, these results indicate IL glutamatergic neurons directly target both excitatory and inhibitory insula neurons.

**Figure 13:**
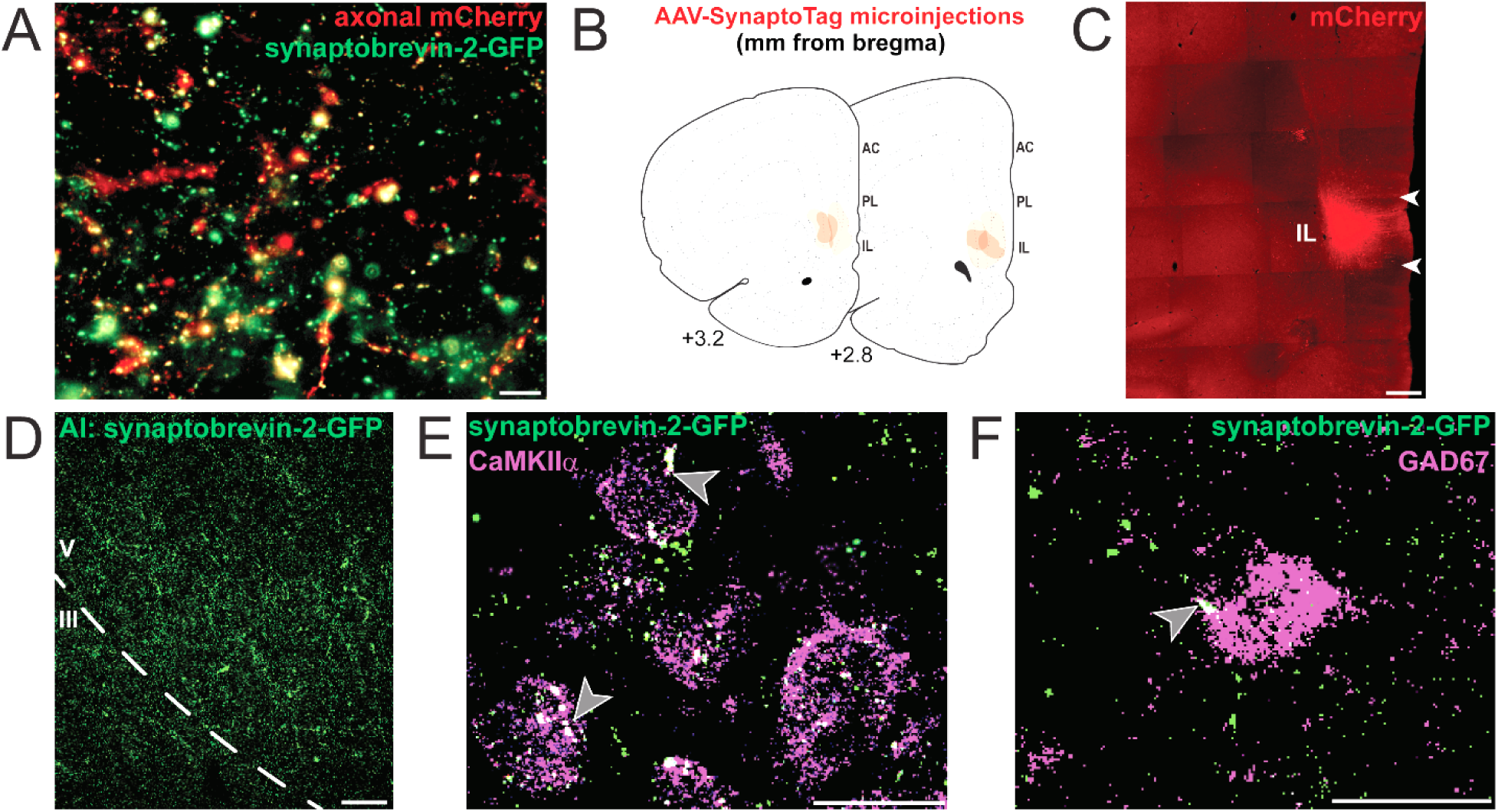
IL projections target AI excitatory and inhibitory neurons. AAV-SynaptoTag utilizes the synapsin promoter to express cell-filing mCherry (labeling soma and axons) and GFP conjugated to synaptobrevin-2 to label pre-synaptic terminals (A), scale bar: 5 μm. Microinjections (n = 4) were mapped onto Swanson Rat Brain Atlas (3^rd^ edition) coronal sections ns (B). A representative injection site (C), scale bar: 500 μm. IL pre-synaptic terminals labeled by GFP in the superficial and deep layers of the AI (D), scale bar: 100 μm. IL pre-synaptic terminals appose CaMKIIα (E) and GAD67 (F) immunoreactive neurons in AI layer VI, scale bar:: 20 μm. Grey arrowheads: representative appositions. AC: anterior cingulate cortex, AI: anterior insular cortex, AAV: adeno-associated virus, CaMKIIα: Ca2+/calmodulin-dependent protein kinase II alpha, GAD67: glutamic acid decarboxylase, 67 kDa isoform, IL: infralimbic cortex, PL: prelimbic cortex.

## 4. Discussion

The current study used a knockdown approach to examine chronic stress effects on long-term neural activity in the frontal lobe, and the role of the IL in coordinating these changes. A lentiviral-packaged construct coding for vGluT1 siRNA was used to knockdown pre-synaptic vesicular glutamate packaging and release (Schuske & Jorgensen, 2004); thus, reducing excitatory outflow from the IL (Myers, et al., 2017). This approach was combined with an anatomical survey of cortical cells expressing ΔFosB, a stable immediate-early gene product and transcription factor that accumulates in response to repeated neural activation (McClung et al., 2004; Nestler, 2015; Nestler et al., 2001, 2002). Across multiple heterogeneous cortical divisions, notable CVS effects included increased ΔFosB in the IL and decreased ΔFosB in the insular cortices. Moreover, decreased ΔFosB expression in the insula was prevented by reduced IL glutamate output. To examine this potential IL-AI circuit, a genetically-encoded marker of IL pre-synaptic terminals was used to label direct synaptic inputs to pyramidal cells and interneurons of the insula. Collectively, these data suggest that IL glutamate signaling mediates excitatory/inhibitory balance in the insula to facilitate cellular responses to chronic stress.

We also used the vGluT1 siRNA approach to determine whether IL output affects coping behaviors or the behavioral consequences of chronic stress. Knockdown of IL glutamate release decreased immobility and increased active coping in the forced swim test, both in stress-naïve and chronically-stressed rats. Additionally, CVS-induced avoidance behaviors in the open field test and novel object interaction were prevented by siRNA treatment. In the elevated plus maze, vGluT1 siRNA decreased risk aversion as evidenced by the increased exploration of the outer ends of open arms. Similar effects on behavioral inhibition were evident in the shock probe assay as siRNA animals spent less time freezing and more time exploring the probe. Importantly, the behavioral effects in siRNA-treated animals were not due to deficits in nociception or associative learning. Taken together, the current data highlight the importance of IL output for facilitating both the neural and behavioral responses to chronic stress.

Multiple studies indicate that local IL circuitry is modified by chronic stress, which shifts excitatory/inhibitory balance and modifies the structure and activity of IL neurons (Cook & Wellman, 2004; Gilabert-Juan et al., 2013; McKlveen et al., 2016, 2019; Radley et al., 2006). Further, chronic stress decreases mPFC gene expression related to GABAergic and glutamatergic signaling, including vGluT1 (Ghosal et al., 2019). IL glutamatergic pyramidal neurons project to multiple forebrain, hypothalamic, and brainstem nuclei involved in stress-integration (Vertes, 2004; Wood et al., 2019). Additionally, recent evidence suggests that IL glutamatergic efferents play a critical role in endocrine and autonomic responses to chronic stress (Myers et al., 2017; Schaeuble et al., 2019). However, IL cortical interactions and their regulation by chronic stress are less clear. The quantitative mapping of ΔFosB+ cells provided an activity-based atlas of chronic stress in the frontal cortex. CVS increased ΔFosB in the IL, an effect that is specific to non-habituating heterotypic stress regimens (Bollinger et al., 2019; Flak et al., 2012; Lehmann & Herkenham, 2011). This was extended to show that ΔFosB expression is specific to pyramidal cells and not observed in interneurons. Other effects of CVS include an increase in the superficial PL, as well as decreases in deep AC and deep VO. Deep AC and VO decreases, and the widespread decreases in the AI and PI, were reversed by siRNA. In addition, vGluT1 siRNA increased ΔFosB immunoreactivity in multiple prefrontal and orbital cell groups after CVS. Ultimately, the data indicate that widespread IL circuit integration in the frontal cortex mediates excitatory/inhibitory activity during chronic stress.

The prefrontal, orbital, and insular regions are implicated in cognitive, emotional, and executive functions (Gianaros & Wager, 2015; Gogolla, 2017; Jahn et al., 2010; McKlveen et al., 2015; Oppenheimer & Cechetto, 2016; Price, 1999). In rodents, prefrontal ΔFosB expression after chronic social defeat associates with social behavior, while PL ΔFosB over-expression reduces social interaction and forced swim activity after chronic defeat (Vialou et al., 2014, 2015). Thus, prefrontal ΔFosB actions, likely as a transcriptional regulator, modulate behavioral responses to social stress. In our study, chronically reduced IL glutamate output altered ΔFosB expression and prevented avoidance behaviors after CVS. In addition, decreased IL output increased active coping and exploratory behaviors. The finding that IL projections are necessary for stress-induced anxiety-like and depression-like behaviors fits with previous work where increased excitability of IL glutamate neurons produced social avoidance and anhedonic behavior (Ferenczi et al., 2016). However, these studies differ from those employing acute mPFC optogenetic stimulation which found increased active coping and reward seeking (Covington et al., 2010; Fuchikami et al., 2015). These divergent results may arise from differences in technical approaches or the specific cell groups targeted. However, numerous behavioral and neurochemical functions in mPFC have been observed to function with an inverted-U response curve (Arnsten, 2007; Bentley et al., 2011; Berridge & Arnsten, 2013; Cools & D’Esposito, 2011; Giustino & Maren, 2018; Luine & Frankfurt, 2012; Sandi & Pinelo-Nava, 2007). This hypothesis suggests that optimal context-specific activity is necessary for neurobehavioral adaptation; consequently, high and low levels of activity could produce similar effects.

Although the function of the IL-AI circuit for behavioral regulation remains to be determined, the insula is involved in action selection based on outcome values (Parkes et al., 2018). The AI also regulates social behavior dependent on the age and stress experience of the interactor (Rogers-Carter et al., 2018), while the PI is necessary for conditioned fear inhibition (Foilb et al., 2016). Collectively, the pivotal role of the insula in behavioral regulation suggests that IL-dependent insula inhibition after chronic stress likely contributes to behavioral outcomes. Interestingly, the insular cortex also responds to visceral sensory information and may link behavior with changes in systemic physiology (Gianaros & Wager, 2015; Oppenheimer & Cechetto, 2016; Yasui et al., 1991). Thus, prefrontal-insular interactions could integrate the central and peripheral responses to chronic stress. This is especially intriguing considering our results with IL vGluT1 knockdown. Here, we show the necessity of IL glutamate output for CVS-induced behaviors as the knockdown reduced behavioral responsiveness to chronic stress. Previously, we found that the knockdown increases physiological responses to CVS. In fact, vGluT1 knockdown increases hypothalamic-pituitary-adrenal responses to acute stress and exacerbates corticosterone responses to a novel stressor after CVS (Myers et al., 2017). Further, IL vGluT1 siRNA and CVS interact to increase cardiovascular stress reactivity, impair vascular function, and promote cardiac and arterial hypertrophy (Schaeuble et al., 2019). Thus, reduced IL output uncouples the positive association between the behavioral and physiological responses to chronic stress, leading to the de-integration of stress responses. Overall, these results highlight the critical role of IL glutamatergic output for coordinating and integrating neural, behavioral, autonomic, and endocrine adaptation to chronic stress.

There are limitations to the current study worth discussion. Similar to previous rodent studies of mPFC and stress processing, these experiments were limited to males. Although, recent studies have demonstrated that chronic stress-induced morphological changes in mPFC pyramidal neurons are similar in male and female rats (Anderson et al., 2019) and that homotypic chronic stress does not affect mPFC ΔFosB expression differentially by sex (Bollinger et al., 2019). However, more widespread frontal lobe changes are sexually-divergent (Carvalho-Netto et al., 2011; Moench et al., 2019) and the behavioral consequences of chronic stress vary by sex (Borrow et al., 2018; Smith et al., 2018). Thus, future studies examining females would provide a better representation of the neural and behavioral consequences of chronic stress. Additionally, given our experimental focus on IL-dependent responses to chronic stress, we did not include a No CVS siRNA group. Although the siRNA may affect basal ΔFosB expression, there were significant effects in the context of chronic stress that indicate a prominent role for the IL. Another consideration is the effect of repeated testing in some of the behavioral assays. Given this design, we cannot isolate the effects of CVS from those due to a second assay exposure. Accordingly, our statistical analysis accounted for this factor as ‘day’ as opposed to CVS. However, treatment-dependent differences within CVS groups indicate that IL glutamate output is critical for the behavioral changes that occurred across testing days during chronic stress exposure. Also, our No CVS GFP controls were reserved for ΔFosB analysis and associative conditioning. Therefore, some behaviors were compared to stress-naïve controls (forced swim, open field, novel object, and conditioning) while others were compared to animals that had already received 1-2 stressors (elevated plus maze, shock probe burying, and Von Frey). However, these tests were carried out 3-12 hours after the most recent stressor which may limit acute stress effects on behavior.

In conclusion, the current study found that IL output was necessary for numerous CVS-induced changes in long-term neural activity, including decreased ΔFosB+ cell density in the insular cortices. The results of CVS and vGluT1 knockdown differed across cortical layers and rostral-caudal gradients of each structure suggesting that regional cell populations differ in functional connectivity and stress-responsiveness. These results were accompanied by circuit mapping that demonstrated direct IL projections to insular pyramidal and inhibitory neurons. Additionally, coping responses and behavioral changes during CVS were dependent on IL output. Taken together, these results emphasize the importance of highly coordinated activity in stress-sensitive frontal lobe networks for both the neural and behavior aspects of chronic stress. Ultimately, determining the mechanisms for excitatory/inhibitory balance in chronic stress-responsive cell groups may identify novel avenues for promoting resilience and adaptation.

## Abbreviations

AC: anterior cingulate cortex
AI: anterior insular cortex
AId: anterior insular cortex, dorsal part
AIv: anterior insular cortex, ventral part
BA25: Brodmann’s area 25
CaMKIIα: Ca^2+^/calmodulin-dependent protein kinase II alpha
CVS: chronic variable stress
EPM: elevated plus maze
FST: forced swim test
GAD67: glutamic acid decarboxylase, 67 kDa isoform
GFP: green fluorescent protein
IL: infralimbic cortex
LO: lateral orbital cortex
LV: lentivirus
MDD: major depressive disorder
MO: medial orbital cortex
mPFC: medial prefrontal cortex
NOI: novel object interaction
OFT: open field test
PI: posterior insular cortex
PL: prelimbic cortex
siRNA: small interfering RNA
SPB: shock probe burying
T: tone only
TS: tone-shock pairing
VF: Von Frey assay
vGluT1: vesicular glutamate transporter 1
VO: ventral orbital cortex

## Acknowledgements

This work was supported by NIH grants R00 HL122454 and R01 HL150559 to Brent Myers and R01 MH049698 to James P. Herman. The authors also thank Sarah Fourman for technical assistance. Jessica M. McKlveen contributed to this article in her personal capacity. The views expressed herein are those of the authors and do not necessarily represent the views of the National Institutes of Health, National Center for Complementary and Integrative Health, or the United States Government. All other authors have no declarations of interest.

## CRediT author statement

**Sebastian A. Pace:** Validation, Formal analysis, Investigation, Data Curation, Writing - Original Draft, Writing - Review & Editing, Visualization

**Connor Christensen:** Validation, Investigation

**Morgan K. Schackmuth:** Validation, Investigation

**Tyler Wallace:** Software, Validation, Investigation, Formal analysis, Data Curation

**Jessica M. McKlveen:** Investigation, Writing - Review & Editing

**Will Beischel:** Investigation

**Rachel Morano:** Investigation

**Jessie R. Scheimann:** Investigation

**Steven P. Wilson:** Resources

**James P. Herman:** Resources, Writing - Review & Editing, Funding acquisition

**Brent Myers:** Conceptualization, Methodology, Formal analysis, Investigation, Writing - Original Draft, Writing - Review & Editing, Visualization, Supervision, Funding acquisition

## Notes

### Competing Interest Statement

The authors have declared no competing interest.

